# Na_V_1.6 inhibition drives the efficacy of voltage-gated sodium channel inhibitors to prevent electrically induced seizures in both wild type and *Scn8a^N1768D/+^* gain-of-function mice

**DOI:** 10.1101/2023.08.03.551823

**Authors:** JP Johnson, Thilo Focken, Parisa Karimi Tari, Celine Dube, Samuel J Goodchild, Jean-Christophe Andrez, Girish Bankar, Kristen Burford, Elaine Chang, Sultan Chowdhury, Jessica Christabel, Richard Dean, Gina de Boer, Christoph Dehnhardt, Wei Gong, Michael Grimwood, Angela Hussainkhel, Qi Jia, Kuldip Khakh, Stephanie Lee, Jenny Li, Sophia Lin, Andrea Lindgren, Verner Lofstrand, Janette Mezeyova, Karen Nelkenbrecher, Noah Gregory Shuart, Luis Sojo, Shaoyi Sun, Matthew Waldbrook, Steven Wesolowski, Michael Wilson, Zhiwei Xie, Alla Zenova, Wei Zhang, Fiona Scott, Alison J Cutts, Robin P Sherrington, Raymond Winquist, Charles J Cohen, James R Empfield

**Affiliations:** Department of In Vitro Biology, Xenon Pharmaceuticals Inc, Burnaby, BC, Canada; Department of Chemistry, Xenon Pharmaceuticals Inc, Burnaby, BC, Canada; Department of In Vivo Biology, Xenon Pharmaceuticals Inc, Burnaby, BC, Canada; Department of Compound Properties, Xenon Pharmaceuticals Inc, Burnaby, BC, Canada; Translational Drug Development, Xenon Pharmaceuticals Inc, Burnaby, BC, Canada; Scientific Affairs, Xenon Pharmaceuticals, Inc, Burnaby, BC, Canada; Executive Team, Xenon Pharmaceuticals Inc, Burnaby, BC V5G 4W8, Canada; Neurocrine Biosciences, San Diego, CA 92130, USA

**Keywords:** Na_V_1.6, Scn8a, anti-seizure medicines, SCN8A-DEE, selective inhibitor

## Abstract

Inhibitors of voltage-gated sodium channels (Na_V_s) are important anti-epileptic drugs, but the contribution of specific channel isoforms is unknown since available inhibitors are nonselective. We created a series of compounds with diverse selectivity profiles enabling block of Na_V_1.6 alone or together with Na_V_1.2. Mice with a heterozygous gain-of-function mutation (N1768D/+) in Scn8a (encoding Na_V_1.6) responded with a tonic-clonic seizure to a mild 6 Hz stimulus that was innocuous to wild-type mice. Pharmacologic inhibition of Na_V_1.6 in *Scn8a^N1768D/+^* mice prevented seizures. Inhibitors were also effective in a direct current maximal electroshock seizure assay in wild-type mice. Na_V_1.6 inhibition correlated with efficacy in both models, even without inhibition of other CNS Na_V_ isoforms. Our data suggest Na_V_1.6 inhibition is a driver of efficacy for Na_V_ inhibitor anti-seizure medicines. Selective Na_V_1.6 inhibitors may provide targeted therapies for human Scn8a developmental and epileptic encephalopathies and better tolerated treatments for idiopathic epilepsies.

**Graphical Abstract:** 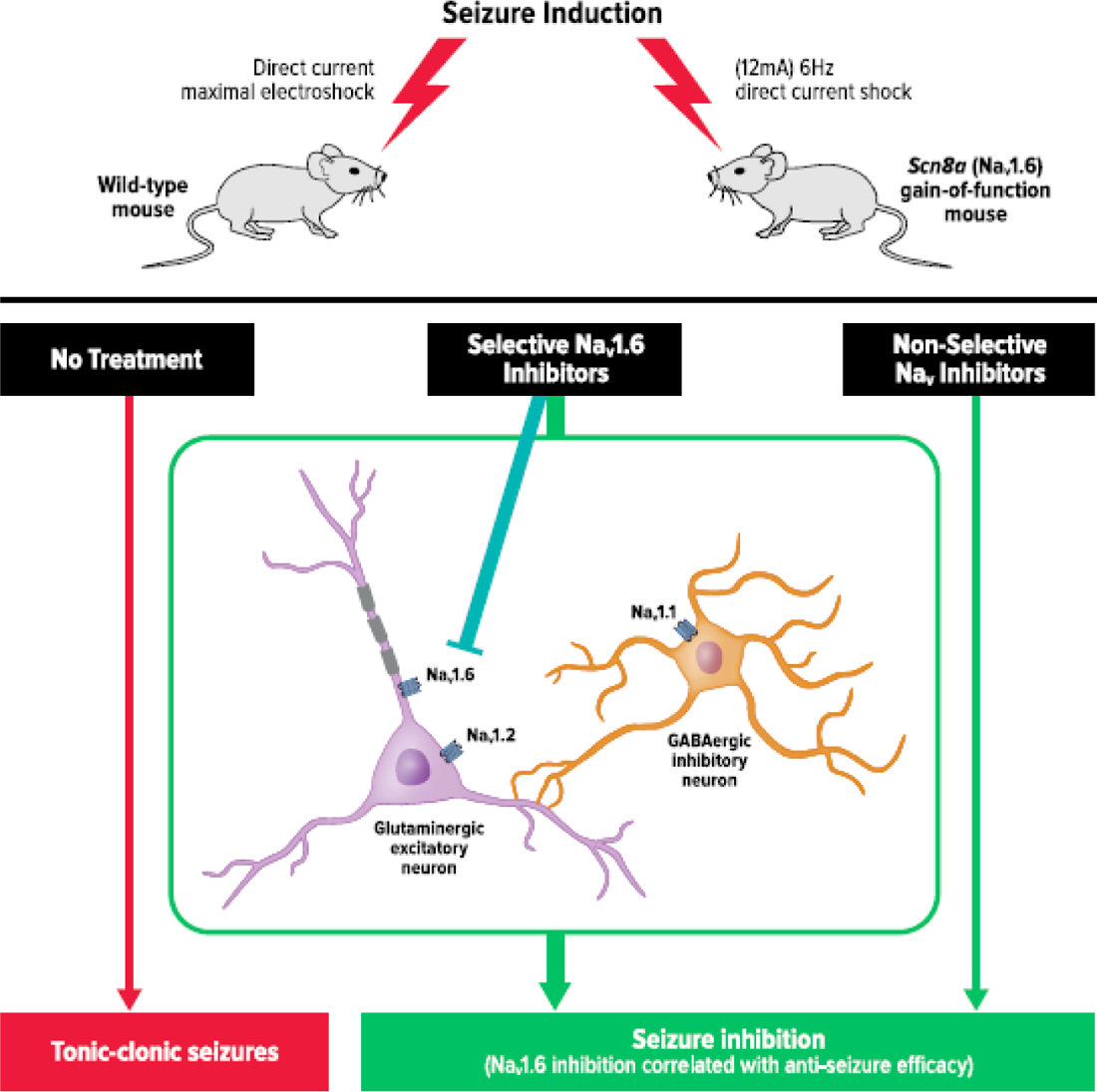

## Introduction

Epilepsy is a common neurologic disorder effecting nearly 4% of people.^1^ Despite dozens of available anti-epileptic drugs (AEDs), more than 30% of epilepsy patients have seizures that are resistant to pharmaceutical treatment.^2^ Uncontrolled seizures, and side effects of currently available drugs, can have dramatic impacts on patients and families both in terms of quality of life and financial security.^3–5^

Central nervous system (CNS) neuron action potential firing is dependent on sodium channels. CNS neurons primarily express four voltage-gated sodium channels (Na_V_s), Na_V_1.1 (encoded by the Scn1a gene), Na_V_1.2 (Scn2a), Na_V_1.3 (Scn3a) and Na_V_1.6 (Scn8a). Pathogenic variants of all four channels have been linked to genetic epilepsies. Na_V_1.3 is primarily expressed embryonically and is not highly expressed in adults except in pathologic conditions. Na_V_1.2 and Na_V_1.6 are both highly expressed in the axon initial segment of excitatory neurons, and gain-of-function (GOF) mutations in both genes have been linked to epilepsy syndromes, SCN2A-DEE and SCN8A-DEE respectively.^6, 7^ In contrast, Na_V_1.1 is a dominant sodium channel in fast firing inhibitory interneurons, and null or loss of function variants of Na_V_1.1 are linked to the severe epileptic encephalopathy known as Dravet syndrome (SCN1A-DEE).^8, 9^ Milder loss of function mutations in Na_V_1.1 cause a less drastic form of epilepsy known as generalized epilepsy with febrile seizures plus (GEFS+).^10^ In both cases, reduced Na_V_1.1 current levels are believed to reduce the firing of inhibitory interneurons, increasing the excitability of neural networks and predisposing patients to seizures.

The human genetic validation of the role of Na_V_s in epilepsy is strengthened by recombinant mouse models with SCN variants. *Scn8a^N1768D/+^*mice are heterozygous for a patient identified GOF variant Na_V_1.6 channel and exhibit reduced seizure thresholds, spontaneous seizures, and early mortality.^7, 11^ Mice with a GOF variant of Scn2a that slows inactivation are also seizure prone.^12^ Mouse models of Dravet Syndrome have been created by heterozygous knock out of Na_V_1.1 and these mice are also prone to seizures and sudden death.^13, 14^

Because Na_V_1.1 is highly expressed in inhibitory interneurons, block of Na_V_1.1 is expected to reduce firing of inhibitory neurons and therefore to be proconvulsant. Available sodium channel inhibitors that block all Na_V_ isoforms with similar potency may be compromised by their activity on Na_V_1.1. Indeed, nonselective Na_V_ inhibitors are contraindicated in Dravet syndrome since they tend to exacerbate seizures.^15, 16^ Patients with GOF variants in Na_V_1.2 and Na_V_1.6 respond well to Na_V_ targeted inhibitors like carbamazepine and phenytoin, and these are first line therapies for those patients.^17, 18^ Nonetheless, nonselective Na_V_ inhibitors have side effects that are likely related to their broad target activity. Nonselective compounds also inhibit the cardiac Na_V_ isoform, Na_V_1.5, and can therefore cause changes in cardiac function. It remains unknown how much of the clinical efficacy of nonselective Na_V_ inhibitors can be attributed explicitly to voltage-gated sodium channels and, more specifically, which sodium channel isoforms are most important for their effectiveness at preventing seizures.

We reasoned that creating new drugs that block Na_V_ channels in excitatory pathways (Na_V_1.6 and Na_V_1.2) while sparing the channels more prominent in inhibitory interneurons (Na_V_1.1) would create a new generation of improved Na_V_ inhibitor drugs. Genetic inhibition of both Nav1.6^19, 20^ and Na_V_1.2^21^ in mice have been shown to reduce seizure susceptibility, lending further support to our hypothesis.

To create selective inhibitors, we targeted a site in the domain IV voltage sensor (VSD4) that has previously proven susceptible to Na_V_1.3 and Na_V_1.7 selective chemistries.^22, 23^ Through an extensive medicinal chemistry effort, we created a set of novel brain penetrant Na_V_1.6 channel inhibitors with diverse selectivity profiles, enabling evaluation of the relative contribution of the CNS Na_V_ isoforms to efficacy in rodent induced seizure models. We evaluated the ability of Na_V_1.6 inhibitors with a range of selectivity profiles to prevent seizures in a modified 6 Hertz psychomotor seizure assay in N1768D/+ mice and in a direct-current maximal electroshock assay (DC-MES) in wild-type mice. Phenytoin and carbamazepine, Na_V_ inhibitors with little selectivity, were compared to novel, selective Na_V_ inhibitors in both mouse seizure models. The results of this study suggest that Na_V_1.6 inhibition is a key driver of seizure control in these mouse models.

## Results

We first set out to create a new class of molecules that can cross the blood brain barrier, specifically target Na_V_1.6 and spare the sodium channels of inhibitory interneurons (Na_V_1.1), and cardiomyocytes (Na_V_1.5). Throughout the course of our work, we created compounds with a broad range brain to plasma ratios and selectivity profiles. **Figure S1** shows the structures of 5 novel aryl sulfonamide compounds with a range of potency and selectivity profiles (XPC-7198, XPC-5462, XPC-7224, XPC-4509, and XPC-6591). The new XPC compounds are expected to bind to the “up” conformation of the domain IV voltage sensor based on their structural similarity to compounds known to bind that region of Na_V_1.3 and Na_V_1.7.^22, 23^ The structures of the commonly prescribed nonselective Na_V_ inhibitors phenytoin and carbamazepine, which bind to the well-conserved internal pore region of Na_V_ channels, are shown for comparison.

Additional selectivity data for XPC-4562 and XPC-7224 in more Na_V_ subtypes and in multiple voltage protocols are described in a separate publication.^24^ Goodchild et al. includes mutagenesis studies to confirm the XPC binding site and further clarifies the functional effects of the XPC Na_V_1.6 inhibitors on channel gating.^24^

### In vitro potency for inhibition of Navs

The adult CNS human Na_V_ isoforms, hNa_V_1.6, hNa_V_1.2, and hNa_V_1.1, as well as for the cardiac isoform hNa_V_1.5 were heterologously expressed in HEK-293 cells and the potency of all 7 test compounds was determined by automated patch-clamp techniques holding the membrane voltage at approximately the V_1/2_ of inactivation for each subtype. **Figure 1** shows concentration-inhibition curves for all 7 compounds shown in **Figure S1**. Phenytoin and carbamazepine are modestly potent and have little isoform selectivity (**Figure 1**). The inhibitor concentrations 50% (IC_50_) of the Na_V_ current for phenytoin were between 3.8 and 9.0 µM for all examined isoforms. Carbamazepine IC_50_ values ranged from 27 to 40 µM, for all isoforms. A summary of fitted IC_50_ values, confidence intervals, and the number of cells tested are shown in **Table S1**.

**Figure 1:**
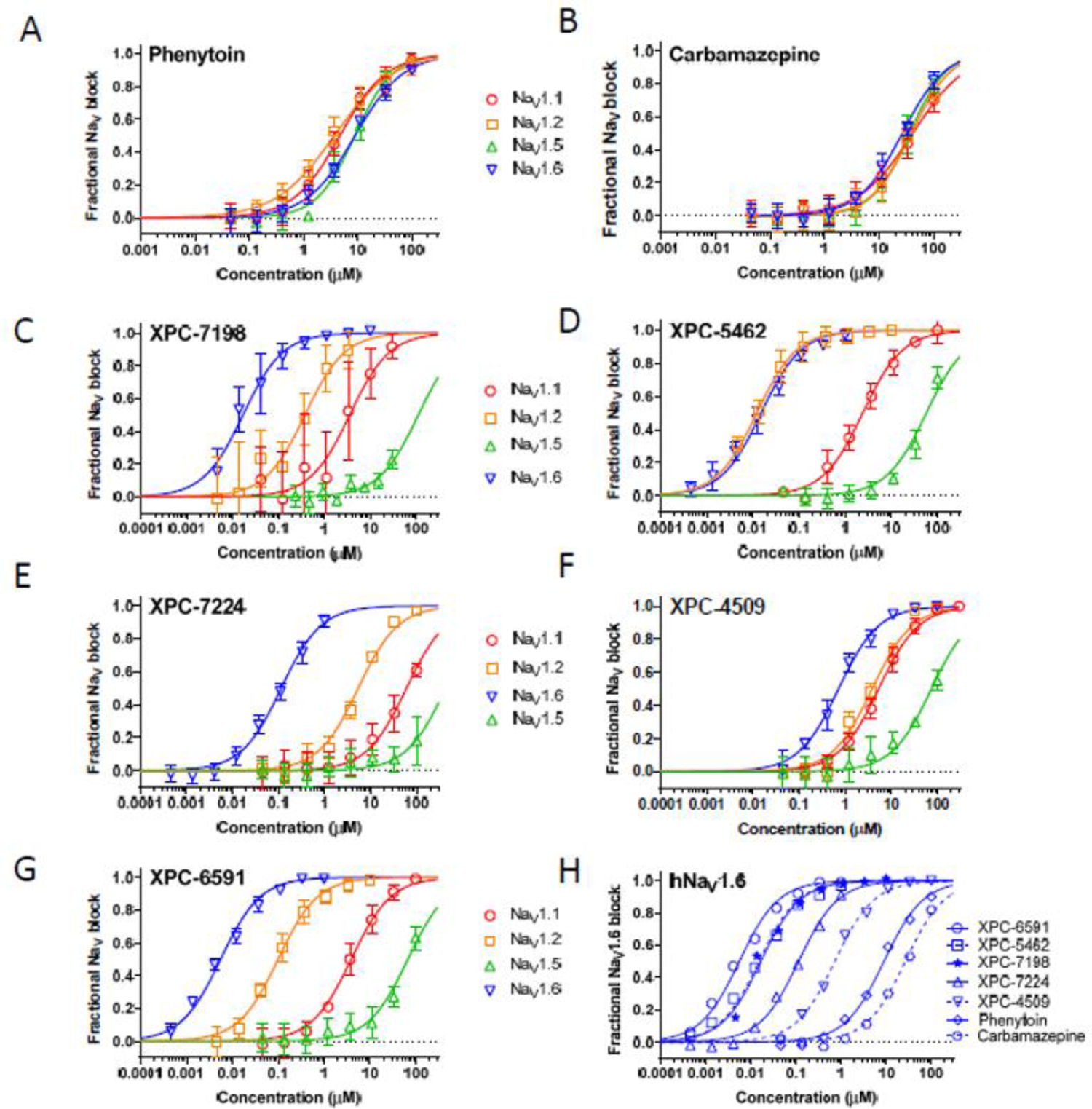
Potency of NaV inhibitor compounds for the adult human CNS and Cardiac NaV channels as determined by automated patch clamp electrophysiology. Phenytoin (A) and carbamazepine (B) have modest potency and little selectivity amongst isoforms. XPC-7198 (C), XPC-5462 (D), XPC-7224, (E) XPC-4509 (F), and XPC-6591 (G) are novel NaV inhibitors with diverse potency and selectivity profiles that were created at Xenon Pharmaceuticals and licensed by Neurocrine Biosciences. (H) shows the relative potency of all 7 compounds on human NaV1.6 with XPC-6591 being the most potent, followed by XPC-7198 > XPC-5462 > XPC-7224 > XPC-4509 >phenytoin > carbamazepine. See Table S1 for fitted IC50 values. Error bars indicate the standard deviations.

Our novel XPC compounds all inhibited Na_V_1.6 more potently than phenytoin or carbamazepine, with a rank order of XPC-6591 < XPC-5462 < XPC-7198 < XPC-7224 < XPC-4509 < phenytoin < carbamazepine. The XPCs displayed a range of potencies (**Figure 1** and **Table S1**). The most potent, XPC-6591, inhibited on hNa_V_1.6 with an IC_50_ of 0.0056 µM. The least potent, XPC-4509, inhibited hNa_V_1.6 with an IC_50_ 0.73 µM.

The binding pocket is well conserved for between human and mouse for Na_V_1.1, Na_V_1.2 and Na_V_1.6, but since the in vivo and ex vivo efficacy experiments used mouse models we also tested the potency of all the compounds on the mouse Na_V_1.6. As expected, potency in mouse NaV1.6 was similar to that for human Na_V_1.6 (**Figure 1** and **Figure S2**).

### Selectivity amongst hNa_V_ isoforms

Phenytoin and carbamazepine inhibited all isoforms with similar potency, consistent with their reputation as nonselective Na_V_ inhibitors (**Figure 2** and **Table S2**). Selectivity ratios for all compounds were generated by dividing the IC_50_ for the compared isoform by the IC_50_ for hNa_V_1.6 (selectivity ratio = IC_50_ hNa_V_1.X / IC_50_ hNa_V_1.6).

**Figure 2:**
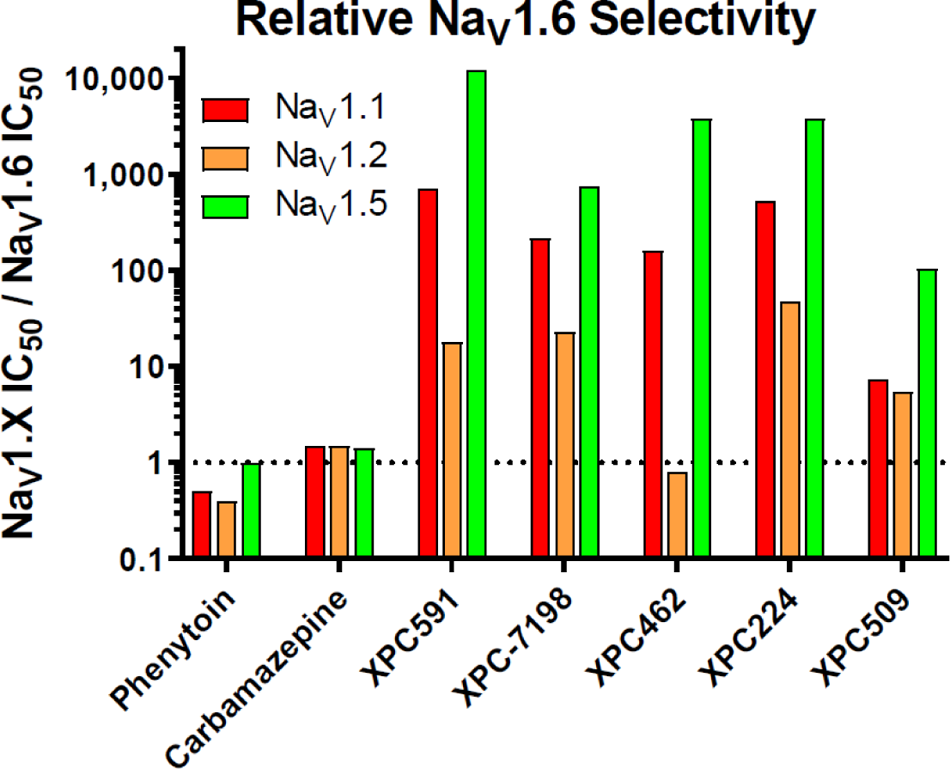
Relative selectivity of NaV inhibitor compounds amongst the NaV channel isoforms highly expressed in adult human CNS and cardiac tissues. Potency values were taken from the dataset in Figure 1. All values are expressed as a ratio of the IC50 on the target channel divided by IC50 on hNaV1.6. Phenytoin and carbamazepine are not selective, while the other compounds have diverse selectivity profiles. See Table S2 for numeric ratios.

Our novel XPC compounds were more selective inhibitors of Na_V_1.6 than phenytoin or carbamazepine but displayed a range of selectivity profiles (**Figure 2** and **Table S2**). All the XPCs were >100-fold selective for Na_V_1.6 vs Na_V_1.5. Selectivity versus Na_V_1.1 ranged from 7-fold (XPC-4509) to >700-fold (XPC-6591). XPC-5462 is slightly more potent on Na_V_1.2 than Na_V_1.6 (ratio 0.8-fold), and the other XPCs ranged up to 47-fold selective for Na_V_1.6 relative to Na_V_1.2. The distinct isoform activities exhibited by this group of seven inhibitors provided an opportunity to query the physiologic impact of distinct selectivity profiles.

Potency and selectivity were also measured in a fluorescent Na^+^ flux assay (**Figure 1** and **Figure S3**). Potency and selectivity measured in the fluorescent assay qualitatively agreed with those measure by electrophysiologic methods.

### Na_V_ inhibitors prevented electrically induced seizures in *Scn8a^N1768D/+^* mice

N1768D was the first SCN8A-DEE variant to be identified in a patient.^7^ N1768D channels open at more negative potentials than wild-type channels, and N1768D also destabilizes Na_V_1.6 inactivation. This leads to persistent currents, excess sodium influx, and neuronal hyperexcitability. Mice bearing an N1768D/+ variant are seizure prone.^11^ Since an excess of Na_V_1.6 current is the underlying cause of the seizure phenotype in patients with SCN8A-DEE, and in the *Scn8a^N1768D/+^* mice, Na_V_ inhibitors are a logical intervention.

We evaluated the test compounds for the ability to inhibit electrically induced seizures in *Scn8a^N1768D/+^* mice. Low intensity (12 mA) 6 Hz Direct current (DC) shocks are innocuous to wild-type mice but littermates heterozygous for N1768D reliably respond to the same intensity shocks with generalized tonic-clonic seizures with hindlimb extension.^25^ We pretreated *Scn8a^N1768D/+^* mice with test compounds two hours prior to the assay and assessed their susceptibility to shock induced seizures (**Figure 3**).

**Figure 3:**
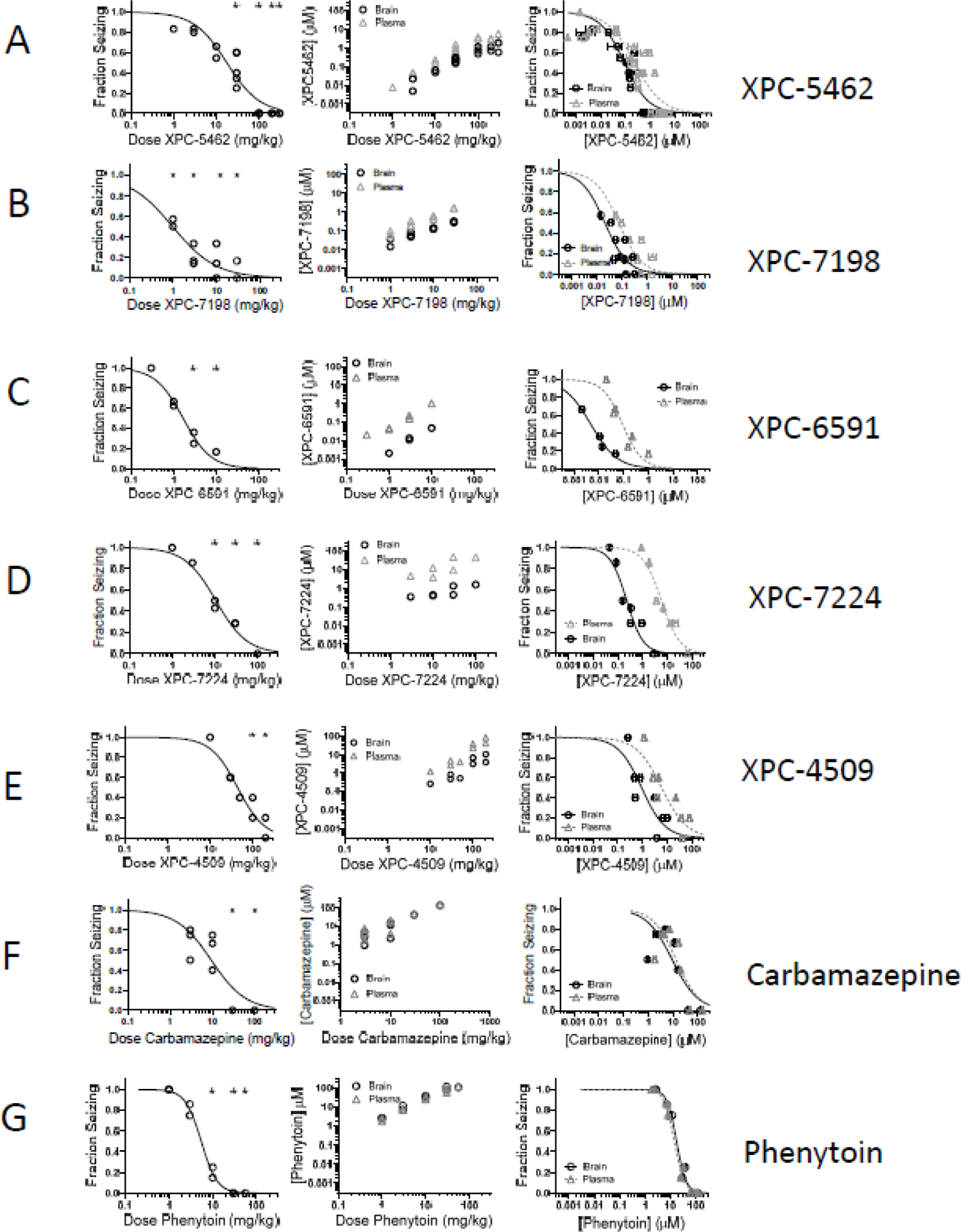
Efficacy in the Scn8a N1768D mouse 6 Hz DC electrical shock assay. The leftmost column shows the fraction of animals that demonstrated a tonic-clonic seizure with hindlimb extension after stimulus as a function of the amount of each compound administered by gastric gavage (or IP injection for phenytoin). Each symbol represents the fraction for a dose group of 6−8 mice. The center column illustrates the brain (black circles) and plasma (gray triangles) concentrations of the tested compounds in the same animals immediately after the efficacy testing. The rightmost column shows the same efficacy data from the left column but plotted versus the mean concentration of each compound in the brain (black circles) or plasma (gray triangles) of the tested animals measured immediately after efficacy testing. Each symbol represents the fraction seizing and mean concentration for a dose group of 6−8 mice. Horizontal error bars indicate the standard error of the mean concentration. All compounds (A) XPC-5462, (B) XPC-7198, (C) XPC-6591, (D), XPC-7224, (E) XPC-4509, (F), carbamazepine, and (G) phenytoin are displayed with identical axes for ease of comparison. For all fits the top of the curve was constrained to 1, the bottom was constrained to 0, and the slope was constrained to <0. See **Table S3** for Fitted ED50 and EC50 values.

All the tested Na_V_ inhibitors effectively protected against seizures in this *Scn8a^N1768D/+^* 6 Hz seizure assay. We evaluated the dose required and the plasma and brain concentrations needed to protect animals from electrical seizure induction.

**Figure 3** shows the dose and concentration dependence of the compounds’ activity in the *Scn8a^N1768D/+^* 6 Hz seizure assay for XPC compounds, carbamazepine, and phenytoin. Doses that provided 50% seizure protection (ED_50_s) in the *Scn8a^N1768D/+^* 6 Hz seizure assay were 18 mg/kg (XPC-5462), 0.98 mg/kg (XPC-7198), 1.7 mg/kg (XPC-6591), 11 mg/kg (XPC-7224), 44 mg/kg (XPC-4509), 9.4 mg/kg (carbamazepine), and 5.5 mg/kg (phenytoin). ED_50_s and confidence intervals are shown in **Table S3**. Statistical analysis is shown in **Figure S4**.

Plasma samples were taken immediately after efficacy testing and the concentrations of the compounds were determined by mass spectrometry. The plasma concentrations associated with 50% efficacy (EC_50_) in the *Scn8a^N1768D/+^* 6 Hz seizure assay were 0.24 µM (XPC-5462), 0.071 µM (XPC-7198), 0.24 µM (XPC-6591), 5.7 µM (XPC-7224), 6.1 µM (XPC-4509), 14 µM (carbamazepine), and 14 µM (phenytoin).

We expect that brain concentrations should be the primary driver of efficacy for CNS indications like epilepsy. Brain tissue samples were taken from all mice immediately after efficacy testing. The brain tissue concentrations of the compounds were measured by mass spectrometry. Brain concentrations that provided 50% effective (EC_50_) protection from seizures were 0.10 µM (XPC-5462), 0.02 µM (XPC-7198), 0.0047 µM (XPC-6591), 0.23 µM (XPC-7224), 0.94 µM (XPC-4509), 9.4 µM (carbamazepine), and 18 µM (phenytoin).

### Comparing in vitro Na_V_1.6 potency and in vivo potency in *Scn8a^N1768D/+^* mice

The plasma concentration versus efficacy relationships for all compounds in the *Scn8a^N1768D/+^* 6 Hz seizure assay are compared in **Figure 4A** and show that compounds were effective at a range of plasma concentrations, an effect can be partially explained by the potency of the compounds. Plotting brain concentration versus efficacy demonstrated that compounds more potent on Na_V_1.6 are effective at lower brain concentrations. The rank order of the brain EC_50_s of compounds in the in vivo assay (XPC-6591 < XPC-7198 < XPC-5462 < XPC-7224 < XPC-4509 < carbamazepine < phenytoin) closely paralleled the potency of the compounds for inhibiting Na_V_1.6 in vitro (XPC-6591 < XPC-7198 < XPC-5462 < XPC-7224 < XPC-4509 < phenytoin < carbamazepine). Carbamazepine performed slightly better than phenytoin in the assay despite being somewhat less potent on Na_V_1.6. This difference may be due to assay variability or may imply additional efficacy for carbamazepine mediated by other targets than Na_V_1.6.

**Figure 4:**
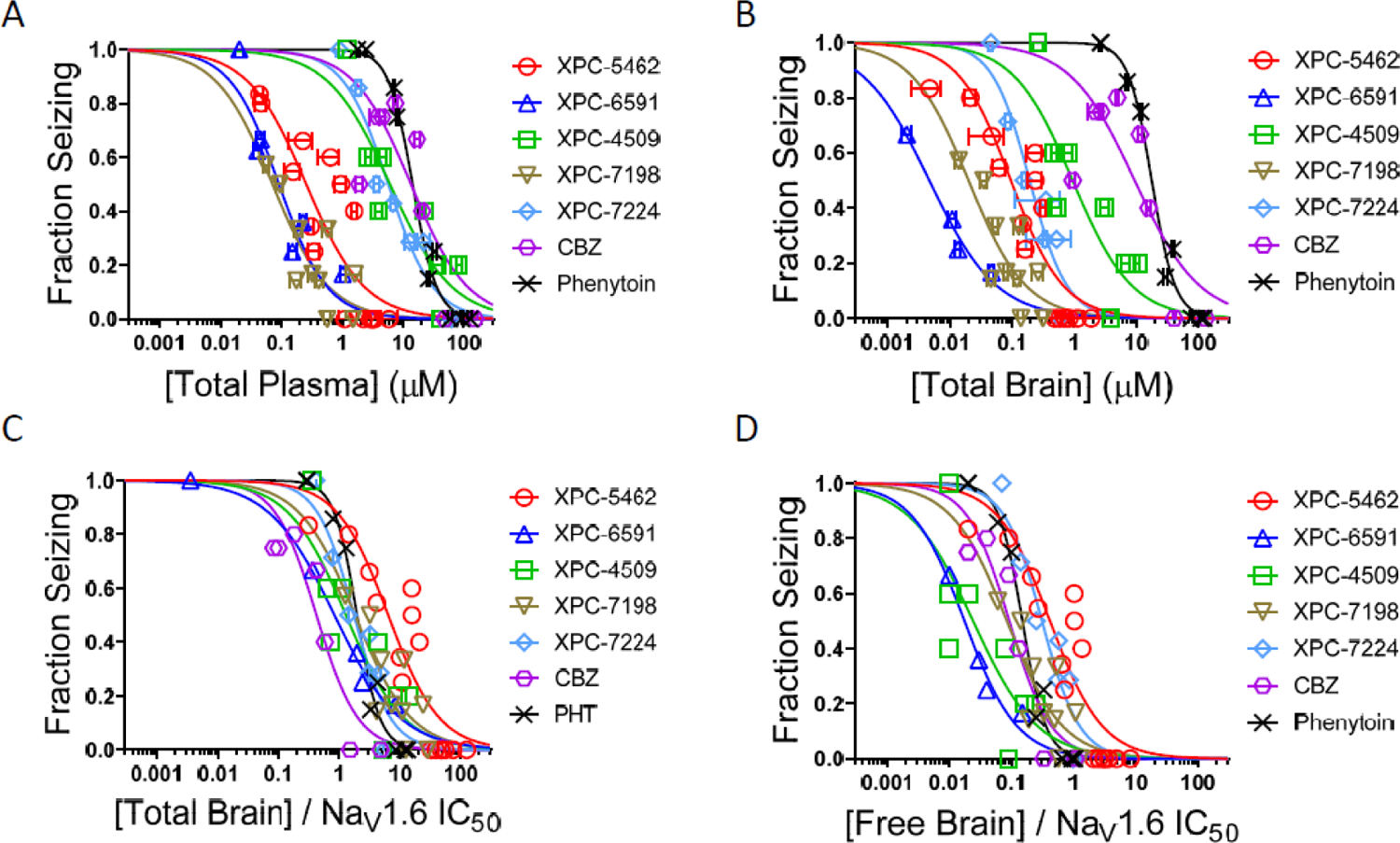
Comparison of Scn8a 6 Hz seizure assay efficacy for all compounds plotted as a function of measured plasma or brain concentrations. Fraction of animals that demonstrated a tonic-clonic seizure with hindlimb extension after stimulus as a function of the amount of each compound administered by gastric gavage (or IP injection for phenytoin) in (A) plasma and (B) total brain. Each symbol represents the fraction for a dose group of 6–8 mice. All compounds are displayed with identical axes for ease of comparison. For all fits the top of the curve was constrained to 1, the bottom was constrained to 0, and the slope was constrained to <0. (C) The fraction seizing in A was normalized for compound potency on NaV1.6 (mean concentrations / NaV1.6 IC50). (D) in the fraction seizing in C was normalized for the brain free fraction (Total Brain Concentration * Brain Homogenate Free Fraction / NaV1.6 IC50).

Accounting for both brain concentration and potency by dividing in vivo brain EC_50_ by the in vitro Na_V_1.6 IC_50_ aligns the curves closely together (**Figure 4C**), illustrating the correlation between in vivo and in vitro activity.

Correcting for free brain concentration of the drugs, scaling the fraction of compound bound in brain homogenate binding assays (**Figure S5**), provided a similar, though noisier, in vivo-in vitro correlation relative to using total brain concentrations (**Figure 4C** and **4D**).

### Na_V_ inhibitors prevented electrically induced seizures in wild-type mice

It is intuitive that inhibition of Na_V_1.6 currents should reduce seizures in a mouse model that is seizure prone due to excess Na_V_1.6 current. But the questions remains whether the same would be true in wild-type mice that are not sensitized by an excess of Na_V_1.6 currents. To address this, we tested all the compounds in an electrically induced seizure model in wild-type mice, the direct current maximal electroshock assay (DC-MES).^25^

**Figure 5** and **Table S4** show the dose and concentration dependence of the compounds’ activity in the DC-MES seizure assay for XPC-5462 (A), XPC-7198 (B), XPC-6591 (C), XPC-224 (D), XPC-4509 I, carbamazepine (F), and phenytoin (G). ED_50_ doses in the DC-MES seizure assay were 23 mg/kg (XPC-5462), 0.80 mg/kg (XPC-7198), 1.5 mg/kg (XPC-6591), 3 mg/kg (XPC-7224), 62 mg/kg (XPC-4509), 25 mg/kg (carbamazepine), and 4.5 mg/kg (phenytoin). Confidence intervals are shown in **Table S4**. Statistical analysis is shown in **Figure S6**.

**Figure 5:**
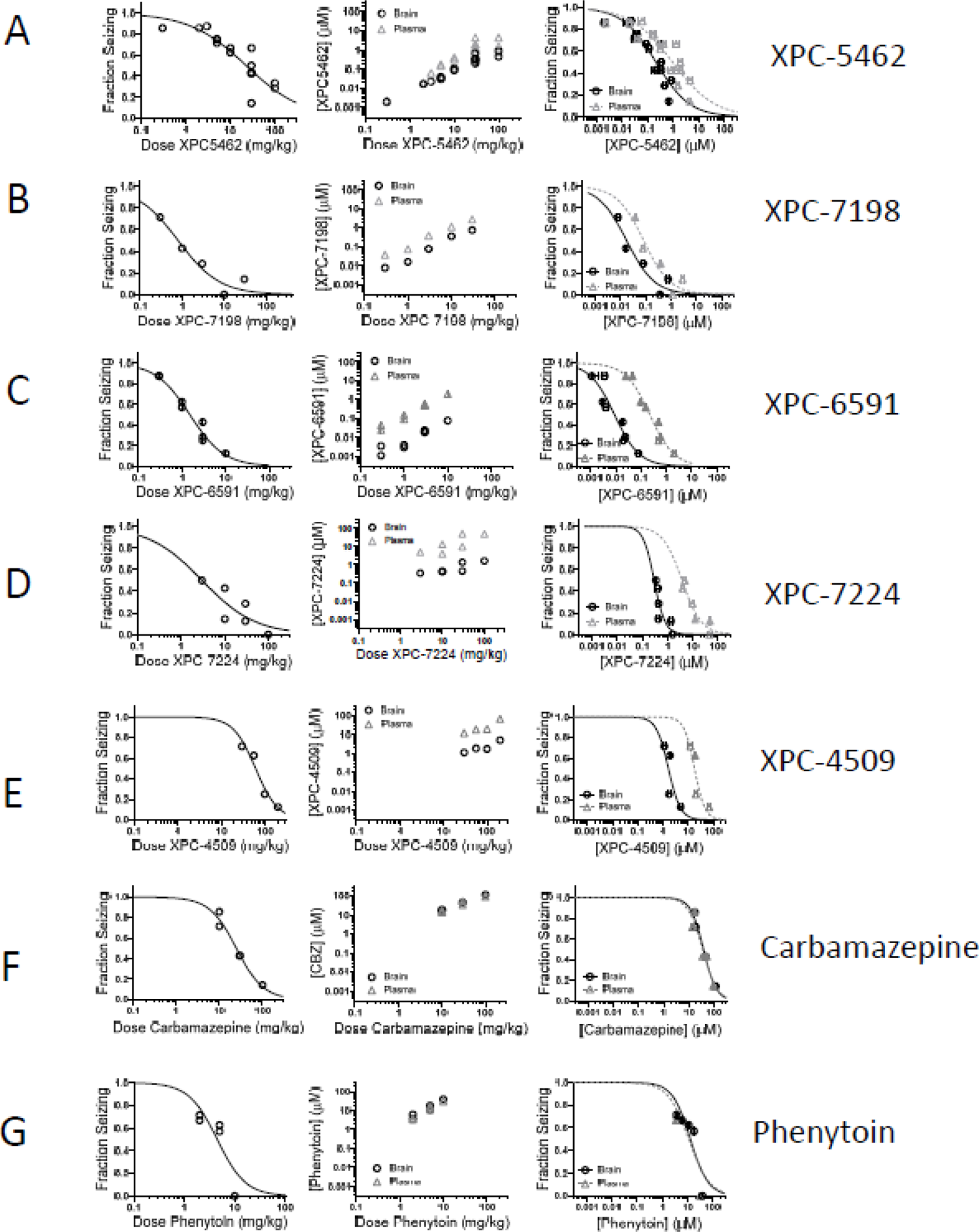
Efficacy in the WT mouse DC-MES electrical shock assay. The leftmost column shows the fraction of animals that demonstrated a tonic-clonic seizure with hindlimb extension after stimulus as a function of the amount of each compound administered by gastric gavage (or IP injection for phenytoin). Each symbol represents the fraction for a dose group of 6–8 mice. The center column illustrates the brain (black circles) and plasma (gray triangles) concentrations of the tested compounds in the same animals immediately after the efficacy testing. The rightmost column shows the same efficacy data from the left column plotted versus the mean concentration of each compound in the brain (black circles) or plasma (gray triangles) of the tested animals measured immediately after efficacy testing. Each symbol represents the fraction seizing and mean concentration for a dose group of 6−8 mice. Horizontal error bars indicate the standard error of the mean concentration. All compounds (A) XPC-5462, (B), XPC-7198, (C) XPC-6591, (D), XPC-7224, (E) XPC-4509, (F), carbamazepine, and (G) phenytoin are displayed with identical axes for ease of comparison. For all fits the top of the curve was constrained to 1, the bottom was constrained to 0, and the slope was constrained to <0. See **Table S3** for Fitted ED50 and EC50 values.

Plasma samples were taken immediately after efficacy testing and the concentrations of the compounds were determined by mass spectrometry. The plasma concentrations associated with 50% efficacy (EC_50_) in the DC-MES seizure assay were 0.93 µM (XPC-5462), 0.081 µM (XPC-7198), 0.22 µM (XPC-6591), 3.8 µM (XPC-7224), 18 µM (XPC-4509), 34 µM (carbamazepine), and 11 µM (phenytoin).

Brain tissue samples were taken from all mice immediately after efficacy testing. The brain tissue concentrations of the compounds were measured by mass spectrometry. Brain concentrations that provided 50% effective (EC_50_) protection from seizures were 0.21 µM (XPC-5462), 0.02 µM (XPC-7198), 0.01 µM (XPC-6591), 0.29 µM (XPC-7224), 1.6 µM (XPC-4509), 38 µM (carbamazepine), and 14 µM (phenytoin).

### Comparing in vitro Na_V_1.6 potency and in vivo potency in wild-type mice

The plasma concentration versus efficacy relationships for all compounds in the DC-MES seizure assay are compared in **Figure 6A** and show that compounds were effective at a range of plasma concentrations, an effect can be partially explained by the potency of the compounds. Plotting brain concentration versus efficacy clarified that compounds more potent on Na_V_1.6 are effective at lower brain concentrations. The rank order of the brain EC_50_s of compounds in the in vivo DC-MES assay (XPC-6591 < XPC-7198 < XPC-5462 < XPC-7224 < XPC-4509 < phenytoin < carbamazepine) was the same as that for the potency of the compounds for inhibiting Na_V_1.6 in vitro (XPC-6591 < XPC-7198 < XPC-5462 < XPC-7224 < XPC-4509 < phenytoin < carbamazepine).

**Figure 6:**
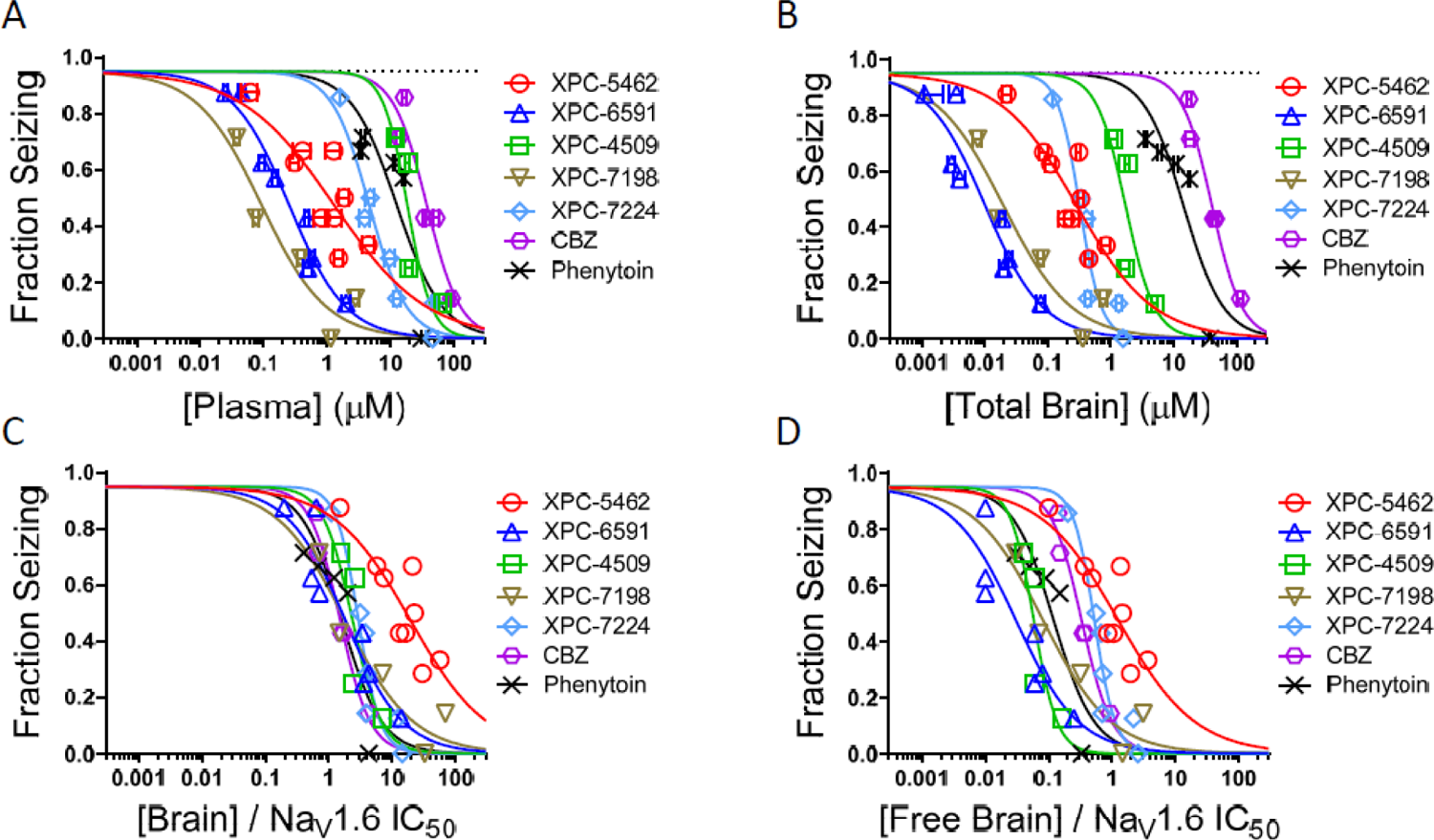
Comparison of WT DC-MES seizure assay efficacy for all compounds plotted as a function of measured plasma or brain concentrations. The fraction of animals that demonstrated a tonic-clonic seizure with hindlimb extension after stimulus as a function of the amount of each compound administered by gastric gavage (or IP injection for phenytoin) for plasma (A) and brain (B) concentrations. (C) Fraction seizing in A was normalized for compound potency on NaV1.6 (mean concentrations / NaV1.6 IC50). (D) Fraction seizing in C was normalized for the brain free fraction (Total Brain Concentration * Brain Homogenate Free Fraction / NaV1.6 IC50).

Accounting for both brain concentration and potency by dividing in vivo brain EC_50_ by the in vitro Na_V_1.6 IC_50_ aligns the curves closely together (**Figure 6C**), illustrating the correlation between in vivo and in vitro activity.

Correcting for free brain concentration of the drugs, by scaling by the fraction of compound bound in brain homogenate binding assays (**Figure S5**), made the in vivo-in vitro correlation worse relative to using total brain concentrations suggesting total brain concentration may be driving Na_V_1.6 target engagement (**Figure 6C** and **6D**).

**Figure 7A** demonstrates the close correlation between efficacy in the two distinct induced seizure models. **Figure 7B** shows the relationship between Na_V_1.2 and Na_V_1.6 potency in vitro and in vivo efficacy in the Scn8a model. **Figure 7C** shows the alignment of in vitro potency and in vivo efficacy in the WT DC-MES model. In both models, compounds that are more potent on Na_V_1.2 and Na_V_1.6 are more potent in vivo. The potency for Na_V_1.6 is most closely and directly aligned with concentrations efficacious in vivo.

**Figure 7:**
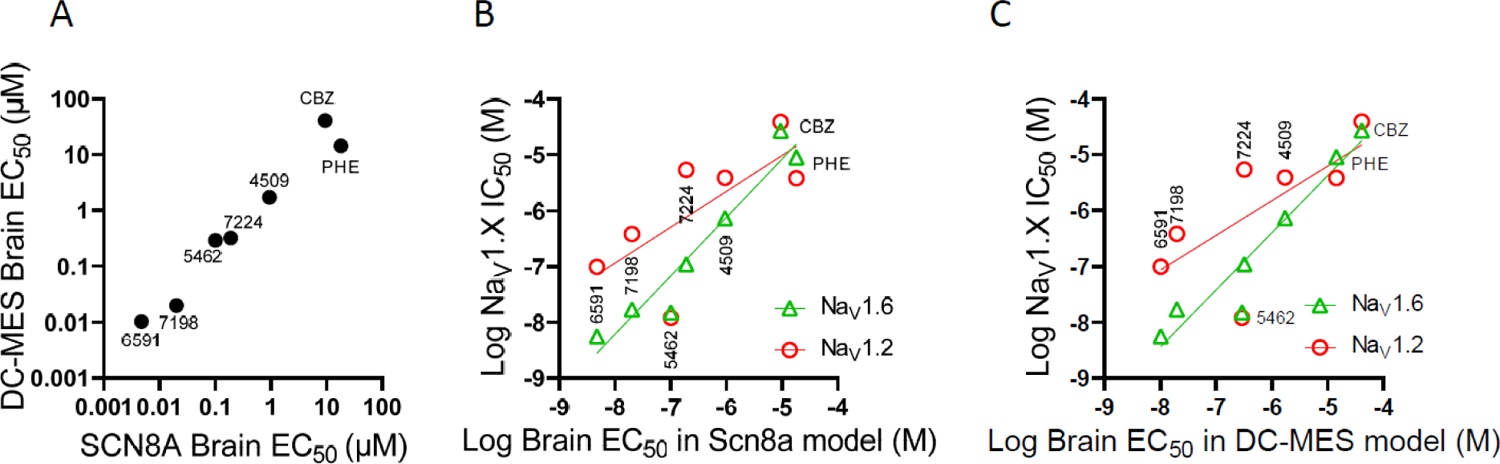
Relative potency in Scn8a mice versus WT mice. (A) Comparison of efficacy in Scn8a mouse 6 Hz seizure assay and the wild type mouse DC-MES assay. (B) 50% effective brain concentrations (Brain EC50) in the Scn8a 6 Hz assay compared to in vitro IC50 for inhibition of hNaV1.2 and hNaV1.6. (C) 50% effective brain concentrations (Brain EC50) in the Scn8a 6 Hz assay compared to in vitro IC50 for inhibition of hNaV1.2 and hNaV1.6.

## Discussion

Sodium channel inhibitors are widely used in clinical practice.^26^ For some, sodium channel inhibition is believed to be the primary driver of efficacy, especially for epilepsy drugs like carbamazepine and phenytoin, and for type I antiarrhythmics, like quinidine and procainamide. But thus far the sodium channel inhibitors available for clinical use do not distinguish amongst the distinct channel isoforms. The lack of selective inhibitors makes understanding the relative importance of distinct channel isoforms for any given physiologic function challenging. For example, type I antiarrhythmics, intended to target the cardiac channel Na_V_1.5, are frequently dose limited by the CNS side effects since they inhibit CNS isoforms with similar potency as the cardiac isoform. Likewise, anti-seizure medications can lead to cardiac side effects in some patients since they inhibit Na_V_1.5 at the same concentrations as the CNS isoforms. The cardiac liabilities of Na_V_ inhibitor ASMs are underscored by the recent US FDA warning that lamotrigine can cause heart rhythm problems.^27^ The FDA also suggested that these cardiac risks should be considered for all pan Na_V_ inhibitor ASMs and listed 10 other drugs that will also require further post market studies to better asses their cardiac liabilities.

Within the CNS, genetics suggests that block of either Na_V_1.6 or Na_V_1.2 would protect against seizures since gain-of-function variants of both isoforms lead to epilepsy syndromes. But inhibition of Na_V_1.1 should be avoided since Na_V_1.1 inhibition is expected to promote seizures. This is supported by compelling genetic evidence that loss of function variants of SCN1A cause GEFS+ and Dravet syndrome. Use of nonselective Na_V_ inhibitors is broadly contraindicated in Dravet Syndrome, since they are known to exacerbate seizures.^28, 29^ The problems of nonselective Na_V_ inhibition in Dravet Syndrome extends beyond seizures; treatment of young Dravet patients with these drugs leads to poorer cognitive development outcomes.^30^

We have been working to address the issues of nonselective Na_V_ inhibitors by developing selective inhibitors of the Na_V_s of excitatory CNS neurons, Na_V_1.6 and Na_V_1.2. NBI-921352 is a selective Na_V_1.6 inhibitor that was created at Xenon Pharmaceuticals and is being developed by Neurocrine Biosciences.^25^ NBI-921352 recently entered human trials to assess its efficacy in SCN8A-DEE patients and in focal onset seizure patients.^31, 32^ During the effort to discover selective inhibitors, like NBI-921352, we created diverse compounds with a range of potency and selectivity profiles. We have made use of some of these compounds with distinct profiles to query the distinct contributions of different channel isoforms to anti-seizure efficacy and neural hyperexcitability.

We compared the relative efficacy of Na_V_ inhibitors with a broad range of potencies for Na_V_1.6 (0.0056 µM for XPC6591 to 27 µM for carbamazepine). We found that in vivo potency in the MES assay was well correlated with the potency for in vitro inhibition of Na_V_1.6.

These inhibitors also displayed a range of selectivity profiles. XPC-7224 is highly selective for Na_V_1.6; it is 47X less potent on Na_V_1.2 and even less potent on other isoforms. XPC-5462 inhibits Na_V_1.6 and Na_V_1.2 with near identical potencies but remains highly selective against the dominant subtype found in inhibitory interneurons (Na_V_1.1) and that of cardiac myocytes (Na_V_1.5). Phenytoin and carbamazepine display nearly equivalent potencies for all the Na_V_ isoforms tested while XPC-7198 and XPC-4509 have intermediate selectivity profiles.

Despite the differences in channel potency and selectivity, all the tested Na_V_ inhibitors were effective at reducing the susceptibility of mice to electrically induced seizures. This was true in Scn8a GOF N1768D/+ mice, and in wild-type mice. The in vitro potency of the compounds was proportional to the in vivo efficacy in both mouse seizure models once the ability of the compounds to get into the brain was accounted for (**Figure 4C**, **Figure 6C**). In addition, the effective brain concentrations were similar in the *Scn8a^N1768D/+^* mouse assay and the wild type mouse assay (**Figure 7A**) for all these compounds.

We have found that measuring tissue concentrations is critical for comparing compounds, since doses do not translate well across species and compounds can have varying levels of CNS penetration. Additionally, the same dose of a given compound can result in very different plasma and tissue concentrations if the precise conditions of the experiment are not the same. The strain of mice, age of mice, dosing method, vehicle composition, method of drug formulation, and even the individual experimenter formulating and administering the compound can all impact the ultimate concentrations achieved in the mouse tissue. Even when all factors are controlled as tightly as possible, mouse tissue concentrations can vary significantly from experiment to experiment. For these reasons we routinely quantitate the plasma concentrations of mice in most of our studies, and we often also measure the levels in the brain or other relevant tissue as well.

Free drug concentrations are often thought to better predict in vivo activity than total drug concentrations, but we have not found that to be the case for this project (**Figures 4** and **6**). We propose two likely possible explanations for this disconnect. First, Na_V_1.6 is directly embedded in the neuronal plasma membrane. The binding site for these compounds, at the domain IV VSD, is directly in contact with the membrane lipid.^24^ Thus the “bound” compound, including that bound to the lipid surrounding the channel binding site, may have direct access to bind the channel. Local anesthetic Na_V_ inhibitors have long been known to access sodium channels via either the aqueous intracellular space or via an intramembrane route, by a mechanism referred to as the modulated receptor hypothesis.^33, 34^ A similar scenario has been suggested by other authors to explain the pharmacology of adrenergic receptors.^35, 36^

Another possibility is that the poorer correlation of the free brain concentration to activity is an artifact of adding another experimentally measured value (the brain homogenate free fraction) to the correlation. Multiplying the brain concentration by the experimentally determined free brain ratio would increase the noise/error of data and hence might degrade the apparent correlation.

Inhibition of Na_V_1.2 may not be as critical for efficacy in these animal models. The potency on Na_V_1.6 appeared to better predict in vivo efficacy and the correlation between in vitro Na_V_1.2 potency and in vivo efficacy was not as strong (R^2^ = 0.51 and 0.49 for *Scn8a^N1768D/+^* mice and wild-type mice respectively) as that for Na_V_1.6 (**Figure 7**; R^2^ = 0.93 and 0.92 for *Scn8a^N1768D/+^* mice and wild-type mice respectively). Despite this, Na_V_1.2 potency does follow the same trend as in vivo efficacy and we cannot rule out a contribution of Na_V_1.2 to seizure prevention.

While these in vivo experiments did not clearly implicate Na_V_1.2 as a driver of improved efficacy, we have found that in ex vivo brain slices seizure models, seizure-like events were differentially impacted by Na_V_1.6 selective XPC-7224 versus dual Na_V_1.2 and Na_V_1.6 inhibitor XPC-5462.^24^ In that scenario XPC-5462, which is equipotent on Na_V_1.2 and Na_V_1.6 was more effective at reducing the incidence of seizure-like activity than the Na_V_1.6 selective XPC-7224. This suggests that there are likely to be some circumstances where inhibiting both Na_V_1.2 and Na_V_1.6 offers improved efficacy over blocking either channel alone.

Our intentions for creating more selective Na_V_ inhibitors are to create more efficacious and better tolerated drugs for patients with epilepsy and seizure syndromes. An inherent risk of the selective approach is that multiple Na_V_ isoforms may be involved in the efficacy of pan Na_V_ inhibitors and stripping the activity for some isoforms (Na_V_1.1 and Na_V_1.2) might simultaneously reduce the anti-seizure activity. Our data do not support this hypothesis and instead suggest that selective Na_V_1.6 inhibitors retain the ability to prevent seizures and that selectivity is not a disadvantage for efficacy, at least in the scenario of the two tested in vivo mouse models.

We have not tested these compounds in mice with Scn2a gain-of-function variants. It might be expected that in such mice, Na_V_1.2 activity would be more important for efficacy, and there may be other models or epilepsy indications where Na_V_1.2 inhibition could be advantageous.

We expect that the group of tool compounds presented here will prove useful for dissecting the relative contributions of different Na_V_ isoforms in neuronal function and physiology. An additional manuscript describes the biophysical mechanisms of Na_V_1.6 inhibition by XPC-4562 and XPC-7224 as representative compounds of this series. This should further facilitate and inform the best use of these compounds for physiologic investigation.^24^

In summary, the in vivo efficacy data presented here supports that selective inhibitors of Na_V_1.6 may be efficacious in human patients, both those with known SCN8A-DEE and in idiopathic epilepsy patients without known variants of Scn8a. We anticipate that there may be patients, or clinical indications, where Na_V_1.6 inhibition is not sufficient for maximal seizure suppression, and in such cases a more pleiotropic compound that inhibits other Na_V_ isoforms might be preferred. In patients where Na_V_1.6 selective inhibitors are sufficient for efficacy, our preclinical studies suggest that they will be better tolerated than currently available pan Na_V_ inhibitors.^25^

## Materials and Methods

### Cell lines

HEK293 cells either stably transfected or transiently transfected. All the cell lines tested negative for mycoplasma. The cell lines were authenticated by STR profiling at EuroFins Genomics. Stable cell lines were transfected with an expression vector containing the full-length cDNA coding for specific human and mouse sodium channel α-subunit, grown in culture media containing 10% fetal bovine serum, and 0.5 mg/mL Geneticin (G418) at 37°C with 5% CO_2_. The Na_V_1.x stable cell lines and accessory constructs used correspond to the following GenBank accession numbers: Human Na_V_1.1 (NM_006920); mouse Na_V_1.6 (NM_001077499); human Na_V_1.2 (NM_021007); human Na_V_1.5 (NM_198056); human Na_V_1.6 (NM_014191); The human Na_V_ β1 subunit (NM_199037) was co-expressed in all cell lines. Na_V_1.6 channels were also co-expressed with human FHF2B (NM_033642) to increase functional expression. Human Na_V_1.2 channels were also co-expressed with Contactin 1 (NM_001843) to increase functional expression.

### Animals

After delivery, animals were allowed sufficient time to acclimate prior to testing (∼1 week). All animals were housed in plastic cages in rooms with controlled humidity, ventilation, and lighting (12 hr/12 hr light–dark cycle). All animal procedures were performed using protocols approved by Xenon Animal Care Committee and the Canadian Council on Animal Care.

### Scn8aN1768D^/+^ mice

Xenon Pharmaceuticals Inc. licensed the mouse with the missense mutation p.Asn1768Asp (N1768D) in the neuronal sodium channel Na_V_1.6, characterized and developed by Dr. M Meisler (University Of Michigan, MI, USA). The Scn8aN1768D knock-in allele was generated by TALEN targeting of (C57BL/6JXSJL) F2 eggs at the University of Michigan Transgenic Animal Model Core. The line was propagated by backcrossing N1768D/+ heterozygotes to C57BL/6J wild-type mice (The Jackson Laboratory, Bar Harbor, ME). Male N1768D/+ heterozygotes on a C57BL/6J background were subsequently backcrossed to C3HeB/FeJ female mice. All the experiments were performed using animals following at least 7 such backcrosses. Experiments were performed using (B6 × C3He) F7 (F7. N1768D/+) offspring aged 35–42 days.

### WT mice

Adult male CF-1 WT albino mice 26–35 g were obtained from Charles River, Senneville, Quebec, Canada. All the assays were carried out in mice 9 to 12 weeks of age.

#### Electrophysiological determination of potency and selectivity

### Current inhibition

Electrophysiology experiments were performed with HEK293 cells. Data was collected using the Qube 384 (Sophion) automated voltage-clamp platform using single hole plates. We used a “V_1/2_” assay where the membrane was held at a voltage close to the empirically determined V_1/2_ for inactivation ^24^. The holding potentials were: −68 mV for Na_V_1.6, Na_V_1.1 and Na_V_1.2, and –100 mV for Na_V_1.5. To assess inhibition, we applied a 0.04 Hz repeating test pulse to −20 mV for 20 ms preceded by a brief 2 ms recovery pulse to −120 mV that was included to increase stability of the currents over time. The protocol was run in vehicle conditions at 0.1 Hz for 150 seconds compound to establish the baseline for each cell prior to the addition of compound for 10 min at a pulse frequency of 0.04 Hz. Full inhibition was defined by adding tetrodotoxin (TTX, 300 nM), or tetracaine for Na_V_1.5 (10 µM), to each well at the end of the experiment. Compounds were then exposed at a single concentration for 10 minutes. One-sixth of every experimental plate was dedicated to vehicle-only wells that enabled correction for nonspecific drift (i.e., rundown) of the signal in each experiment. For all channel subtypes, inhibition by the compound reached steady state within 10 minutes of incubation. The current inhibition values (I(CPD)) were normalized to both the vehicle (Icontrol) and the full response defined by supramaximal TTX (ITTX) or tetracaine (for Na_V_1.5) addition responses according to Equation 1:

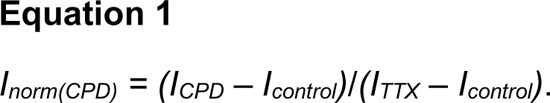

This normalized inhibition was then further normalized to the span of the assay to account for the run-down seen in cells exposed to vehicle alone for 20 minutes as follows:

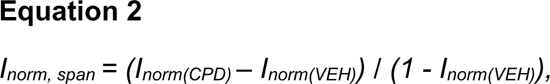

where:

*I _norm, span_* = the current response normalized to within the span of the assay.

*I_norm(CPD)_* = the normalized response in the presence of compound.

*I_norm(VEH)_* = the normalized response in the absence of compound.

This normalization ensures that the data ranges were between 0 and 1, and there is no rundown in the plots. The normalized data from all cell recordings at a concentration were grouped together and plotted with GraphPad Prism 8, and IC_50_ values were calculated for grouped data using the following version of the Hill equation:

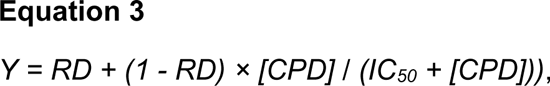

where:

*Y* = the fraction of sodium current blocked in the presence of the compound.

*[CPD]* = the concentration of compound.

*IC_50_* = the IC_50_ concentration.

*RD* = the “rundown” of sodium current in vehicle alone, which is equal to 0 in this case, as the inhibition has already been normalized to the span. The Hill slope was fixed to 1. The 95% CI for the IC_50_ from the fitted curve to the mean data were reported unless otherwise noted.

All Qube experiments were performed at 27°C ± 2°C.

### Automated patch-clamp recording solutions

The recording solutions contained: Intracellular solution (ICS): 5 mM NaCl, 10 mM CsCl, 120 mM CsF, 0.1 mM CaCl_2_, 2 mM MgCl_2_, 10 mM HEPES (4-(2-hydroxyethyl)-1-piperazineethanesulfonic acid buffer), 10 mM EGTA (ethylene glycol tetraacetic acid); adjusted to pH 7.2 with CsOH. Extracellular solution (ECS): 140 mM NaCl, 5 mM KCl, 2 mM CaCl_2_, 1 mM MgCl_2_, 10 mM HEPES; adjusted to pH 7.4 with NaOH. Osmolarity in all ICS and ECS solutions was adjusted with glucose to 300 mOsm/kg and 310 mOsm/kg, respectively.

### In Vitro Sodium Influx Assays

Our sodium influx assays employ the use of the cell permeable, sodium sensitive dye ANG2 to quantify sodium ion influx through sodium channels which are maintained in an open state by use of sodium channel modulators. These high throughput sodium influx assays allow for rapid profiling and characterization of sodium channel blockers. The specific modulators and concentrations used to keep each Na_V_ subtype in an open conformation are shown below.

|  | 1.6 | 1.5 | 1.2 | 1.1 |
| --- | --- | --- | --- | --- |
| **Nav**<br>**Modulator(s)** | 25 μM<br>Veratridine,<br>50 nM ATX-II | 500 nM<br>Veratridine,<br>100 nM ApC | 6 μM<br>Veratridine,<br>30 nM ATX-II | 5 μM<br>Veratridine,<br>50 nM ATX-II |

Trex HEK293 cells were stably transfected with an inducible expression vector containing the full-length cDNA coding for the desired human sodium channel β-subunit and with an expression vector containing full length cDNA coding for the β1-subunit or FHF2B. Sodium channel expressing cell lines were induced with tetracycline (1μg/mL) and plated on 384-well PDL- coated plates at a density of 25K-30K cells/ well in culture media (DMEM, containing 10% FBS and 1% L-glutamine). After overnight incubation (37°C, 5% CO_2_), culture media was removed and cells were loaded with 5uM ANG2 dye for 1-1.5h in Buffer 1 (155 mM NMDG, 5 mM KCl, 2 mM CaCl_2_, 1 mM MgCl_2_, 10 mM HEPES, 10 mM glucose, adjusted with Tris to pH 7.4). Excess dye was removed, and cells were incubated with and without test compound for 1 hr in buffer 1 containing sodium channel modulator(s) at room temperature. A Hamamatsu FDSS µCell was used to perform a 1:1 addition of Na/K challenge buffer (140 mM NaCl, 20 mM HEPES, 1 mM CaCl_2_, 15 mM KCl, 1 mM MgCl_2_, 10 mM glucose, adjusted with Tris to pH 7.4) and simultaneously read plates at excitation wavelength of 530 nm and emission wavelength of 558 nm. Non-sodium channel mediated sodium influx was determined in the presence of either 1 or 25 µM TTX. This background signal was subtracted from the total sodium influx signal in the absence of compound, all data in the presence of compound was normalized to this control. Percent inhibition of sodium ion influx was calculated for each test compound at each test concentration to determine the IC_50_ values.

### Na^+^ Flux Assay Data Analysis

Percent inhibition in sodium influx assays was determined and IC_50_ values were calculated using a 4-parameter logistic model (% inhibition = (A+((B-A)/(1+((x/C)^D)))) using IDBS XLfit™, where A and B are the maximal and minimum inhibition respectively, C is the IC_50_ concentration, and D is the (Hill) slope.

### Formulation and oral dosing

#### Vehicle preparation

The vehicle for oral dosing solutions was 0.5% methyl cellulose and 0.2% Tween-80 in deionized (DI) water. DI water (0.8 L) was heated up to 70°C to 80°C. Five grams of methyl cellulose was slowly added to heated DI water. The mixture was stirred until it formed a homogeneous milky suspension. The suspension was moved to a cold room and stirred overnight to get a clear solution. Two milliliters of Tween-80 was added to the clear solution and diluted up to 1 L with DI water. The vehicle solution was stored at 2°C to 8°C.

#### Drug formulation

XPC Compounds were weighed into vials. An appropriate amount of vehicle was added to the compound powder and then mixed on a T18 ULTRA TURRAX homogenizer (IKA, Wilmington, NC) to create a uniform suspension at the desired concentration. The vials were then wrapped in aluminum foil to protect them from light and placed on a stir plate until the time of dosing. Carbamazepine was formulated in the same manner. Phenytoin was formulated in 0.9% physiological saline.

### Dosing

All compounds except phenytoin were administered orally using a stainless-steel gavage needle at a dose volume of 10 ml/kg. Phenytoin was formulated in physiologic saline and was administered intraperitoneally using a 25-gauge needle at a dose volume of 10 mL/kg. All compounds were administered 2 hours prior to electrical seizure induction for all seizure models employed in this study.

#### Bioanalytical assessment of plasma and brain concentrations

Sample collection: Approximately 0.5 mL of blood was collected from each mouse at the end of the assay via cardiac puncture under deep anesthesia. The blood samples were collected in a syringe and transferred to tubes containing EDTA. Blood was stored at 4°C until centrifuged within 30 minutes of collection. Plasma was harvested and placed on dry ice and stored in a freezer set to maintain a temperature of −70°C to −80°C until analysis. Brains were harvested immediately after blood collection and placed on dry ice prior to storage in a freezer set to maintain a temperature of −70°C to −80°C until analysis.

Plasma samples: Extraction of plasma samples was carried out by protein precipitation using acetonitrile. Plasma samples (50 µL) were mixed with 50 µL of internal standard (IS) solution in water followed by addition of 10 µL of concentrated ortho-phosphoric acid and 200 µL of acetonitrile. Samples were vortexed for 30 seconds, centrifuged at 13,000 rpm for 20 minutes, decanted in to a 96-well plate, and further centrifuged at 4,000 rpm for 20 minutes. The samples were analyzed by UHPLC-ESI-MS/MS as described below.

Brain samples: Prior to extraction, pre-weighed whole brains were homogenized in 1:1 acetonitrile/water (v/v) (4 mL per mouse brain) using an IKA T18 ULTRA-TURRAX Homogenizer at the setting of 4 for approximately 2 min. The homogenate was centrifuged at 13,000 rpm for 20 min and 50 µL of the supernatant were treated exactly as described above for plasma samples. 50 µL of the brain homogenate were then treated exactly as the plasma samples described above.

Standards and quality control (QC) samples: K2EDTA Blank mouse plasma purchased from Valley Biomedical, California, USA was used to prepare standards and QC samples for plasma quantitation and as surrogates for brain homogenate quantitation. Calibration samples ranged from 2.34 ng/mL to 4,800 ng/mL. QC samples concentration included 14 ng/mL (QC-L), 255 ng/mL (QC-M) and 3,600 ng/mL (QC-H). Standards and QC samples were processed the same way as the sample extracts described above.

Analytical methods and statistics for plasma and brain tissue samples: Samples were analyzed by UHPLC-ESI MS/MS using a Sciex TQ-5500 triple quadrupole mass spectrometer equipped with a Shimadzu Nexera UHPLC pump and auto-sampler system using an ACE C18 PFP, 2.50 x 50 mm, 1.7 µ particle size column and gradient elution consisting of solvent A (0.1% formic acid in water) and solvent B (0.1% formic acid in acetonitrile) starting at 20% B from 0 min to 0.4 min and then increased to 100% B from 0.4 min to 0.6 min. At 2.0 min, the mobile phase composition was switched back to 60% B for 1 min. The flow rate used throughout the experiment was 0.4 min/mL and the column was kept at 40°C. The analytes, XPC-7198, XPC-5462, XPC-7224, XPC-4509, XPC-6591, carbamazepine and phenytoin and their respective internal standards IS1, IS2, IS3, IS4, IS5, IS6, and IS7 were detected by electrospray in the positive ion mode using the following transitions: m/z 492.96/90.9 and m/z 523.10/105 for XPC-7198 and IS1, m/z 487.05/58.1 and m/z 315.1/109.1 for XPC-5462 and IS2, m/z 484.05/105.1 and m/z 443.95/91.1 for XPC-7224 and IS3, m/z 481.16/91.1 and m/z 454.98/148 for XPC-4509 and IS4, m/z 493.99/105 and m/z 514.05/105.1 for XPC-6591 and IS5, m/z 237.0/194 and m/z 488.04/109.1 for carbamazepine and IS6 and m/z 251/42 and m/z 488.04/109.1 for phenytoin and IS7. The UHPLC-ESI MS/MS system was controlled by Analyst 1.6.

Sample concentrations were determined using a linear calibration function, weighted 1/X, generated by the regression of analyte to IS peak area ratios in the standard samples to their respective concentrations. Acceptance criteria for the analytical run required that the back calculated values of the standards and the QC samples fell within ± 20% of their nominal values, except for the lowest standard or lower limit of quantitation (LLOQ), for which the acceptance criterion was ± 25%. At least 6 out of 12 standard points had to show back-calculated values within ± 20% of their nominal concentrations for the calibration to be accepted. At least three QC samples, one at each level, had to show back-calculated values within ± 20% of their nominal concentrations for the whole sample batch to be valid.

### The modified 6 Hz psychomotor seizure assay

Scn8a^N1768D/+^ heterozygous mice were tested at 5 weeks of age (p35). The modified 6 Hz seizure assay in Scn8a^N1768D/+^ heterozygous mice was adapted from the traditional 6 Hz assay psychomotor seizure assay to provide a measure of in vivo on target (Na_V_1.6 mediated) efficacy (Barton et al., 2001). The modified assay used a low frequency (6 Hz) but long-duration stimulation (3 seconds) to induce seizures. We identified 12 mA and a 0.3 millisecond pulse interval as a suitable current for testing in Scn8a^N1768D/+^ mice, since it differentiated mutant and wild-type (WT) mice. An electroshock (6 Hz, 12 mA) was delivered for 3 seconds (at 0.3 millisecond pulse interval) by corneal electrodes (Electro Convulsive Therapy Unit 57800 from Ugo Basile). Immediately prior to the electroshock, the animals’ eyes were anesthetized with a drop of Alcaine (0.5% proparacaine hydrochloride). Upon corneal stimulation, WT mice experienced mild seizure behaviors such as facial clonus, forelimb clonus, Straub tail, rearing, and falling, but did not experience a generalized tonic-clonic seizure (GTC) with hindlimb extension. Scn8a^N1768D/+^ animals, however, in addition to mild seizure behaviors, experienced a GTC with hindlimb extension. The modified assay showed a clear differentiation of seizure behavior between WT and Scn8a^N1768D/+^ mice. Scn8a^N1768D/+^ mice exhibited GTC with hindlimb extension but not WT mice.

Scn8a^N1768D/+^ animals were dosed PO with vehicle or test compound two hours before the administration of the electric stimulation. An animal was considered protected in the assay upon prevention of GTC with hindlimb extension and was then scored “0”. An animal displaying GTC with hindlimb extension was considered not protected and is then scored “1”. The experimenter scoring the seizure behavior was blinded to the treatment.

### DC-Maximal electroshock seizure assay

The maximal electroshock seizure (MES) assay has been extensively used in the search for anticonvulsant substances^37–39^. The MES assay is sensitive to nonselective Na_V_ inhibitors. It is considered a model for generalized tonic-clonic (GTC) seizures and provides an assessment of seizure spread. Briefly, an electroshock of direct current (DC) was delivered by corneal electrodes (Electro Convulsive Therapy Unit 57800 from Ugo Basile). In CF1 mice aged 9 to 12 weeks, a direct current of 50 mA (60 Hz) was delivered for 0.2 seconds (pulse width of 0.5 ms). Immediately prior to the electroshock, the animals’ eyes were anesthetized with a drop of Alcaine (0.5% proparacaine hydrochloride). Upon corneal stimulation, naïve animals experienced a generalized tonic-clonic seizure (GTC) with hindlimb extension.

For the efficacy experiments, single dose and repeated dose, animals were dosed PO with vehicle or test compound two hours before the administration of the electric stimulation. An animal was considered protected in the assay in the absence of a GTC with hindlimb extension and is then scored “0”. An animal displaying GTC with hindlimb extension was considered not protected and is then scored “1”. The experimenter scoring the seizure behavior was blinded to the treatment.

### Blinding of in vivo efficacy experiments

On each testing day, individual treatment groups were assigned a random label (e.g., A, B, C, etc.) by the technical staff administering the compound. To ensure blinding, the technical staff member performing drug administration differed from the person performing the test. Therefore, the experimenter conducting testing was blinded to the treatment group (e.g., drug or vehicle treatment, dose, and time point).

### Randomization of in vivo efficacy experiments

Randomization of animals into various treatment groups occurred on a per-animal (e.g., rather than a per-cage) basis. Therefore, each animal was randomly assigned to a treatment group, and all animals tested in each experiment had an equal chance of assignment to any treatment group. Prior to each study, a randomization sequence was obtained (www.graphpad.com/quickcalcs).

### Quantification and statistical analysis

All statistics were calculated using GraphPad Prism version 8 software. All fraction seizing data plotted by dose groups is expressed and is an absolute fraction of all tested animals that seized (number of animals that seized / the number of animals tested). Between-group differences for the fraction seizing data were compared to vehicle response and were analyzed using a Kruskal-Wallis test followed by Dunn’s multiple comparisons test. Statistical significance was reached at values of *P*<.05. More details of the statistical analysis, with *P* values, are presented in the supplemental figures and legends.

## Acknowledgments

We thank Dr. Miriam Meisler, PhD for making Scn8a^N1768D/+^ mice available. We thank the SCN8A- RES patients, families, physicians, and patient advocates for detailed discussions regarding the clinical presentation of Scn8a variant patients and the unmet medical needs in their community.

## Author Contributions

The experiments in this report were conceptualized by JPJ, TF, PT, C Dube, SG, RD, KK, LS, AC, RS, RW, CC, JE. Experiments were conducted and analyzed by PT, C Dube, SG, EC, JC, GDB, AH, KK, S Lee, JL, S Lin, AL, JM, KN, NS, MW. Cell lines and mutants were created by JM. Medicinal Chemistry expertise was provided by TF, JA, KB, SC, C Dehnhardt, WG, MG, QJ, VL, SS, SW, MW, ZX, AZ, WZ, JE. JPJ, SG, GB, RD, FS, and AC contributed to the writing and editing of the manuscript. All work conducted for this manuscript was under the supervision of JPJ. All authors approved the final manuscript draft for submission.

## Declaration of interests

All Authors, except Fiona Scott, are current or previous employees of Xenon Pharmaceuticals Inc. and may hold equity in that company. Fiona Scott is employed by Neurocrine Biosciences and may hold equity in that company.

## Notes

### Summary of Updates

This manuscript was revised to include the bioRxiv DOI number for reference #24 (Goodchild et al. 2023).

## REFERENCES

1. Hesdorffer, D.C., Logroscino, G., Benn, E.K., Katri, N., Cascino, G., and Hauser, W.A. (2011). Estimating risk for developing epilepsy: a population-based study in Rochester, Minnesota. Neurology 76, 23–27. 10.1212/WNL.0b013e318204a36a.

2. Loscher, W., and Schmidt, D. (2011). Modern antiepileptic drug development has failed to deliver: ways out of the current dilemma. Epilepsia 52, 657–678. 10.1111/j.1528-1167.2011.03024.x.

3. Ronen, G.M., Rosenbaum, P.L., Boyle, M.H., and Streiner, D.L. (2018). Patient-reported quality of life and biopsychosocial health outcomes in pediatric epilepsy: An update for healthcare providers. Epilepsy & behavior: E&B 86, 19–24. 10.1016/j.yebeh.2018.05.009.

4. Baranowski, C.J. (2018). The quality of life of older adults with epilepsy: A systematic review. Seizure 60, 190–197. 10.1016/j.seizure.2018.06.002.

5. Begley, C.E., and Durgin, T.L. (2015). The direct cost of epilepsy in the United States: A systematic review of estimates. Epilepsia 56, 1376–1387. 10.1111/epi.13084.

6. Sugawara, T., Mazaki-Miyazaki, E., Ito, M., Nagafuji, H., Fukuma, G., Mitsudome, A., Wada, K., Kaneko, S., Hirose, S., and Yamakawa, K. (2001). Nav1.1 mutations cause febrile seizures associated with afebrile partial seizures. Neurology 57, 703–705.

7. Veeramah, K.R., O’Brien, J.E., Meisler, M.H., Cheng, X., Dib-Hajj, S.D., Waxman, S.G., Talwar, D., Girirajan, S., Eichler, E.E., Restifo, L.L., et al. (2012). De novo pathogenic SCN8A mutation identified by whole-genome sequencing of a family quartet affected by infantile epileptic encephalopathy and SUDEP. Am J Hum Genet 90, 502–510. 10.1016/j.ajhg.2012.01.006.

8. Claes, L., Del-Favero, J., Ceulemans, B., Lagae, L., Van Broeckhoven, C., and De Jonghe, P. (2001). De novo mutations in the sodium-channel gene SCN1A cause severe myoclonic epilepsy of infancy. Am J Hum Genet 68, 1327–1332. 10.1086/320609.

9. Sugawara, T., Mazaki-Miyazaki, E., Fukushima, K., Shimomura, J., Fujiwara, T., Hamano, S., Inoue, Y., and Yamakawa, K. (2002). Frequent mutations of SCN1A in severe myoclonic epilepsy in infancy. Neurology 58, 1122–1124.

10. Escayg, A., MacDonald, B.T., Meisler, M.H., Baulac, S., Huberfeld, G., An-Gourfinkel, I., Brice, A., LeGuern, E., Moulard, B., Chaigne, D., et al. (2000). Mutations of SCN1A, encoding a neuronal sodium channel, in two families with GEFS+2. Nat Genet 24, 343–345. 10.1038/74159.

11. Wagnon, J.L., Korn, M.J., Parent, R., Tarpey, T.A., Jones, J.M., Hammer, M.F., Murphy, G.G., Parent, J.M., and Meisler, M.H. (2015). Convulsive seizures and SUDEP in a mouse model of SCN8A epileptic encephalopathy. Human molecular genetics 24, 506–515. 10.1093/hmg/ddu470.

12. Kearney, J.A., Plummer, N.W., Smith, M.R., Kapur, J., Cummins, T.R., Waxman, S.G., Goldin, A.L., and Meisler, M.H. (2001). A gain-of-function mutation in the sodium channel gene Scn2a results in seizures and behavioral abnormalities. Neuroscience 102, 307–317.

13. Yu, F.H., Mantegazza, M., Westenbroek, R.E., Robbins, C.A., Kalume, F., Burton, K.A., Spain, W.J., McKnight, G.S., Scheuer, T., and Catterall, W.A. (2006). Reduced sodium current in GABAergic interneurons in a mouse model of severe myoclonic epilepsy in infancy. Nat Neurosci 9, 1142–1149. 10.1038/nn1754.

14. Ogiwara, I., Miyamoto, H., Morita, N., Atapour, N., Mazaki, E., Inoue, I., Takeuchi, T., Itohara, S., Yanagawa, Y., Obata, K., et al. (2007). Nav1.1 localizes to axons of parvalbumin-positive inhibitory interneurons: a circuit basis for epileptic seizures in mice carrying an Scn1a gene mutation. The Journal of neuroscience: the official journal of the Society for Neuroscience 27, 5903–5914. 10.1523/JNEUROSCI.5270-06.2007.

15. Shi, X.Y., Tomonoh, Y., Wang, W.Z., Ishii, A., Higurashi, N., Kurahashi, H., Kaneko, S., Hirose, S., and Epilepsy Genetic Study Group, J. (2016). Efficacy of antiepileptic drugs for the treatment of Dravet syndrome with different genotypes. Brain Dev 38, 40–46. 10.1016/j.braindev.2015.06.008.

16. Genton, P., Gelisse, P., Thomas, P., and Dravet, C. (2000). Do carbamazepine and phenytoin aggravate juvenile myoclonic epilepsy? Neurology 55, 1106–1109. 10.1212/wnl.55.8.1106.

17. Boerma, R.S., Braun, K.P., van de Broek, M.P., van Berkestijn, F.M., Swinkels, M.E., Hagebeuk, E.O., Lindhout, D., van Kempen, M., Boon, M., Nicolai, J., et al. (2016). Remarkable Phenytoin Sensitivity in 4 Children with SCN8A-related Epilepsy: A Molecular Neuropharmacological Approach. Neurotherapeutics: the journal of the American Society for Experimental NeuroTherapeutics 13, 192–197. 10.1007/s13311-015-0372-8.

18. Howell, K.B., McMahon, J.M., Carvill, G.L., Tambunan, D., Mackay, M.T., Rodriguez-Casero, V., Webster, R., Clark, D., Freeman, J.L., Calvert, S., et al. (2015). SCN2A encephalopathy: A major cause of epilepsy of infancy with migrating focal seizures. Neurology 85, 958–966. 10.1212/WNL.0000000000001926.

19. Lenk, G.M., Jafar-Nejad, P., Hill, S.F., Huffman, L.D., Smolen, C.E., Wagnon, J.L., Petit, H., Yu, W., Ziobro, J., Bhatia, K., et al. (2020). Scn8a Antisense Oligonucleotide Is Protective in Mouse Models of SCN8A Encephalopathy and Dravet Syndrome. Ann Neurol 87, 339–346. 10.1002/ana.25676.

20. Martin, M.S., Tang, B., Papale, L.A., Yu, F.H., Catterall, W.A., and Escayg, A. (2007). The voltage-gated sodium channel Scn8a is a genetic modifier of severe myoclonic epilepsy of infancy. Human molecular genetics 16, 2892–2899. 10.1093/hmg/ddm248.

21. Li, M., Jancovski, N., Jafar-Nejad, P., Burbano, L.E., Rollo, B., Richards, K., Drew, L., Sedo, A., Heighway, J., Pachernegg, S., et al. (2021). Antisense oligonucleotide therapy reduces seizures and extends life span in an SCN2A gain-of-function epilepsy model. J Clin Invest 131. 10.1172/JCI152079.

22. McCormack, K., Santos, S., Chapman, M.L., Krafte, D.S., Marron, B.E., West, C.W., Krambis, M.J., Antonio, B.M., Zellmer, S.G., Printzenhoff, D., et al. (2013). Voltage sensor interaction site for selective small molecule inhibitors of voltage-gated sodium channels. Proc Natl Acad Sci U S A 110, E2724–2732. 10.1073/pnas.1220844110.

23. Ahuja, S., Mukund, S., Deng, L., Khakh, K., Chang, E., Ho, H., Shriver, S., Young, C., Lin, S., Johnson, J.P., Jr., et al. (2015). Structural basis of Nav1.7 inhibition by an isoform-selective small-molecule antagonist. Science 350, aac5464. 10.1126/science.aac5464.

24. Goodchild, S.J. (2023). Molecular Pharmacology of Selective Nav1.6 and Dual Nav1.6 & Nav1.2 Channel Inhibitors that Suppress Excitatory Neuronal Activity ex vivo. bioRxiv doi: 10.1101/2023.08.03.551643.

25. Johnson, J.P., Focken, T., Khakh, K., Tari, P.K., Dube, C., Goodchild, S.J., Andrez, J.C., Bankar, G., Bogucki, D., Burford, K., et al. (2022). NBI-921352, a first-in-class, NaV1.6 selective, sodium channel inhibitor that prevents seizures in Scn8a gain-of-function mice, and wild-type mice and rats. Elife 11. 10.7554/eLife.72468.

26. Brodie, M.J. (2017). Sodium Channel Blockers in the Treatment of Epilepsy. CNS Drugs 31, 527–534. 10.1007/s40263-017-0441-0.

27. FDA (2021). Drug Safety Communication: Studies show increased risk of heart rhythm problems with seizure and mental health medicine lamotrigine (Lamictal) in patients with heart disease.

28. Wallace, A., Wirrell, E., and Kenney-Jung, D.L. (2016). Pharmacotherapy for Dravet Syndrome. Paediatr Drugs 18, 197–208. 10.1007/s40272-016-0171-7.

29. Brunklaus, A., Ellis, R., Reavey, E., Forbes, G.H., and Zuberi, S.M. (2012). Prognostic, clinical and demographic features in SCN1A mutation-positive Dravet syndrome. Brain 135, 2329–2336. 10.1093/brain/aws151.

30. de Lange, I.M., Gunning, B., Sonsma, A.C.M., van Gemert, L., van Kempen, M., Verbeek, N.E., Nicolai, J., Knoers, N., Koeleman, B.P.C., and Brilstra, E.H. (2018). Influence of contraindicated medication use on cognitive outcome in Dravet syndrome and age at first afebrile seizure as a clinical predictor in SCN1A-related seizure phenotypes. Epilepsia 59, 1154–1165. 10.1111/epi.14191.

31. Neurocrine. (2021). Study to Evaluate NBI-921352 as Adjunctive Therapy in Subjects With SCN8A Developmental and Epileptic Encephalopathy Syndrome (SCN8A-DEE). ClinicalTrials.gov

32. Neurocrine (2022). Extension Study to Evaluate the Safety and Tolerability of NBI-921352 When Used With Anti-seizure Medications in Adults With Focal Onset Seizures. ClinicalTrials.gov.

33. Korner, J., Albani, S., Sudha Bhagavath Eswaran, V., Roehl, A.B., Rossetti, G., and Lampert, A. (2022). Sodium Channels and Local Anesthetics-Old Friends With New Perspectives. Front Pharmacol 13, 837088. 10.3389/fphar.2022.837088.

34. Hille, B. (1977). Local anesthetics: hydrophilic and hydrophobic pathways for the drug-receptor reaction. J Gen Physiol 69, 497–515. 10.1085/jgp.69.4.497.

35. Sykes, D.A., Parry, C., Reilly, J., Wright, P., Fairhurst, R.A., and Charlton, S.J. (2014). Observed drug-receptor association rates are governed by membrane affinity: the importance of establishing “micro-pharmacokinetic/pharmacodynamic relationships” at the beta2-adrenoceptor. Mol Pharmacol 85, 608–617. 10.1124/mol.113.090209.

36. Dickson, C.J., Hornak, V., Velez-Vega, C., McKay, D.J., Reilly, J., Sandham, D.A., Shaw, D., Fairhurst, R.A., Charlton, S.J., Sykes, D.A., et al. (2016). Uncoupling the Structure-Activity Relationships of beta2 Adrenergic Receptor Ligands from Membrane Binding. J Med Chem 59, 5780–5789. 10.1021/acs.jmedchem.6b00358.

37. Loscher, W., Fassbender, C.P., and Nolting, B. (1991). The role of technical, biological and pharmacological factors in the laboratory evaluation of anticonvulsant drugs. II. Maximal electroshock seizure models. Epilepsy Res 8, 79–94. 10.1016/0920-1211(91)90075-q.

38. Piredda, S.G., Woodhead, J.H., and Swinyard, E.A. (1985). Effect of stimulus intensity on the profile of anticonvulsant activity of phenytoin, ethosuximide and valproate. J Pharmacol Exp Ther 232, 741–745.

39. White, H.S., Johnson, M., Wolf, H.H., and Kupferberg, H.J. (1995). The early identification of anticonvulsant activity: role of the maximal electroshock and subcutaneous pentylenetetrazol seizure models. Ital J Neurol Sci 16, 73–77. 10.1007/BF02229077.

