## Supplemental Materials for "Na_V_1.6 inhibition drives the efficacy of voltage-gated sodium channel inhibitors to prevent electrically induced seizures in both wild type and *Scn8a^N1768D/+^* gain-of-function mice"

### Supplementary Information

#### Supplementary Results

**Table S1** shows the potency of the compounds discussed in the study for the human orthologues of $hNa_V1.1$, $hNa_V1.2$, $hNa_V1.5$, and $hNa_V1.6$. $IC_{50}$ values are determined from the fits shown in **Figure S1**. Ns represent the total number of cells exposed to the test compound. Each cell was exposed to 1 concentration of 1 compound.

**Table S2** shows the selectivity of the compounds discussed in the study for the human orthologues of $hNa_V1.1$, $hNa_V1.2$, $hNa_V1.5$ relative to the potency on $hNa_V1.6$. $IC_{50}$ values are determined from the fits shown in **Figure S1**. Selectivity ratios were determined by dividing the $IC_{50}$ on a given channel by the $IC_{50}$ for $hNa_V1.6$ ($hNa_V1.X$ $IC_{50}$ / $hNa_V1.6$ $IC_{50}$).

**Table S3** The efficacy is compared for all compounds based on the curves fit in **Figure 3.** Efficacy relative to dose is described by the fitted dose providing half maximal efficacy ($ED_{50}$). Efficacy relative to concentration is described by the fitted concentration of half maximal efficacy relative to plasma concentration (Plasma $EC_{50}$), and brain concentration (Brain $EC_{50}$). The final column compares the in vivo potency for inhibition of seizure behaviors to the in vitro potency for inhibition of $Na_V1.6$ current (see **Table S1**).

**Table S4** shows a summary of WT DC-MES assay efficacy data as seen in **Figure 6.**

The efficacy is compared for all compounds based on the curves fit in **Figure S3**. Efficacy relative to dose is described by the fitted dose providing half maximal efficacy ($ED_{50}$). Efficacy relative to concentration is described by the fitted concentration of half maximal efficacy relative to plasma concentration (Plasma $EC_{50}$), and brain concentration (Brain $EC_{50}$). The final column compares the in vivo potency for inhibition of seizure behaviors to the in vitro potency for inhibition of $Na_V1.6$ current (see **Table S1**).

Summary of WT DC-MES assay efficacy data in **Figure S6**.

### Supplementary Information

#### Supplementary Tables

##### Table S1: Potency on $hNa_V1.6$, $hNa_V1.2$, $hNa_V1.1$, and $hNa_V1.5$.

| | | $Na_V1.6$ | $Na_V1.2$ | $Na_V1.1$ | $Na_V1.5$ |
|---|---|---|---|---|---|
| **6591** | $IC_{50}$ (µM) | 0.0056 | 0.099 | 4 | 68 |
| | 95% CI | 0.0049 to 0.0063 | 0.088 to 0.11 | 3.3 to 4. 7 | 52 to 90 |
| | N (cells) | 44 | 46 | 42 | |
| **7198** | $IC_{50}$ | 0.017 | 0.38 | 3.6 | >100 |
| | 95% CI | 0.014 to 0.020 | 0.25 to 0.56 | 2.2 to 6.0 | 92 to 140 |
| | N (cells) | 60 | 38 | 45 | 50 |
| **5462** | $IC_{50}$ (µM) | 0.015 | 0.012 | 2.4 | 57 |
| | 95% CI | 0.013 to 0.018 | 0.011 to 0.014 | 2.0 to 2.8 | 48 to 69 |
| | N (cells) | 53 | 58 | 37 | 27 |
| **7224** | $IC_{50}$ (µM) | 0.11 | 5.4 | 60 | >100 |
| | 95% CI | 0.10 to 0.13 | 4.9 to 6.0 | 49 to 75 | 280 to 760 |
| | N (cells) | 52 | 37 | 39 | 30 |
| **4509** | $IC_{50}$ (µM) | 0.73 | 3.9 | 5.4 | 75 |
| | 95% CI | 0.64 to 0.83 | 3.4 to 4.6 | 5.0 to 5.8 | 56 to 100 |
| | N (cells) | 37 | 45 | 106 | 30 |
| **PHT** | $IC_{50}$ (µM) | 9 | 3.8 | 4.7 | 8.9 |
| | 95% CI | 8.1 to 9.9 | 3.1 to 4.7 | 4.1 to 5.5 | 7.1 to 11 |
| | N (cells) | 67 | 30 | 52 | 21 |
| **CBZ** | $IC_{50}$ (µM) | 27 | 39 | 40 | 36 |
| | 95% CI | 24 to 30 | 33 to 47 | 33 to 48 | 30 to 43 |
| | N (cells) | 59 | 40 | 33 | 44 |

##### Table S2: Selectivity relative to $hNa_V1.6$.

| | 6591 | 7198 | 5462 | 7224 | 4509 | PHT | CBZ |
|---|---|---|---|---|---|---|---|
| **$Na_V1.2$** | 17.9 | 22.5 | 0.8 | 47.1 | 5.4 | 0.4 | 1.5 |
| **$Na_V1.1$** | 709.4 | 213.6 | 158.5 | 526.7 | 7.4 | 0.5 | 1.5 |
| **$Na_V1.5$** | 12224.4 | 752.4 | 3785.2 | 3751.7 | 103.6 | 1.0 | 1.4 |

**Table S3: Comparison of compound efficacy in the SCN8A N1768D 6 Hz seizure assay.**

| | $ED_{50}$ (mg/kg) | 95% CI | Hill Slope | $ED_{50}$ (mg/kg) | 95% CI | Hill Slope | $ED_{50}$ (mg/kg) | 95% CI | Hill Slope | Brain $EC_{50}$ / $Na_V$1.6 $IC_{50}$ |
|---|---|---|---|---|---|---|---|---|---|---|
| **XPC-5462** | 18 | 11 to 26 | -1.1 | 0.24 | 0.11 to 0.39 | -0.9 | 0.10 | 0.044 to 0.14 | -1 | 6.7 |
| **XPC-6591** | 1.7 | 1.2 to 2.5 | -1.3 | 0.24 | 0.045 to 0.24 | -1.1 | 0.0047 | 0.0029 to 0.0068 | -0.8 | 0.84 |
| **XPC-4509** | 44 | 30 to 63 | -1.3 | 6.1 | 2.3 to 15 | -0.9 | 0.94 | 0.27 to 2.5 | -0.9 | 1.3 |
| **XPC-7198** | 0.98 | 0.11 to 1.8 | -0.9 | 0.071 | 0.013 to 0.12 | -0.9 | 0.02 | 0.0032 to 0.036 | -0.9 | 1.8 |
| **XPC-7224** | 11 | 7.5 to 15 | -1.2 | 5.7 | 3.6 to 9.3 | -1.2 | 0.23 | 0.14 to 0.40 | -1.3 | 2.1 |
| **Carbamazepine** | 9.4 | 3.0 to 27 | -1.1 | 14 | 2.6 to 49 | -0.9 | 9.4 | 1.4 to 40 | -0.8 | 0.35 |
| **Phenytoin** | 5.5 | 4.8 to 6.3 | -2.4 | 14 | 12 to 17 | -2.2 | 18 | 14 to 21 | -2.3 | 2 |

**Table S4: Comparison of compound efficacy in the DC-MES assay.**

| | $ED_{50}$ (mg/kg) | 95% CI | Hill Slope | $ED_{50}$ (mg/kg) | 95% CI | Hill Slope | $ED_{50}$ (mg/kg) | 95% CI | Hill Slope | Brain $EC_{50}$ / $Na_V$1.6 $IC_{50}$ |
|---|---|---|---|---|---|---|---|---|---|---|
| **XPC-5462** | 23 | 15 to 37 | -0.7 | 0.93 | 0.56 to 1.6 | -0.6 | 0.21 | 0.14 to 0.32 | -0.7 | 14 |
| **XPC-6591** | 1.5 | 1.2 to 1.9 | -1.1 | 0.22 | 0.16 to 0.30 | -0.9 | 0.01 | 0.0051 to 0.015 | -0.9 | 1.8 |
| **XPC-4509** | 62 | 23 to 110 | -1.7 | 18 | 10 to 32 | -2.3 | 1.6 | 0.74 to 3.5 | -1.9 | 2.2 |
| **XPC-7198** | 0.80 | 0.21 to 1.8 | -0.9 | 0.081 | 0.0082 to 0.24 | -0.9 | 0.02 | 0.0023 to 0.050 | -0.8 | 1.8 |
| **XPC-7224** | 3.0 | ??? to 8.8 | -0.71 | 3.8 | 1.2 to 5.7 | -1.1 | 0.29 | 0.19 to 0.41 | -2 | 2.6 |
| **Carbamazepine** | 25 | 18 to 35 | -1.4 | 34 | 20 to 63 | -1.5 | 38 | 30 to 49 | -1.7 | 1.4 |
| **Phenytoin** | 4.5 | 2.3 to 8.7 | -1.5 | 11 | 3.9 to 31 | -1.1 | 14 | 5.9 to 28 | -1.2 | 1.6 |

**Figure S1: Structures of traditional and novel $Na_V$ inhibitors.**

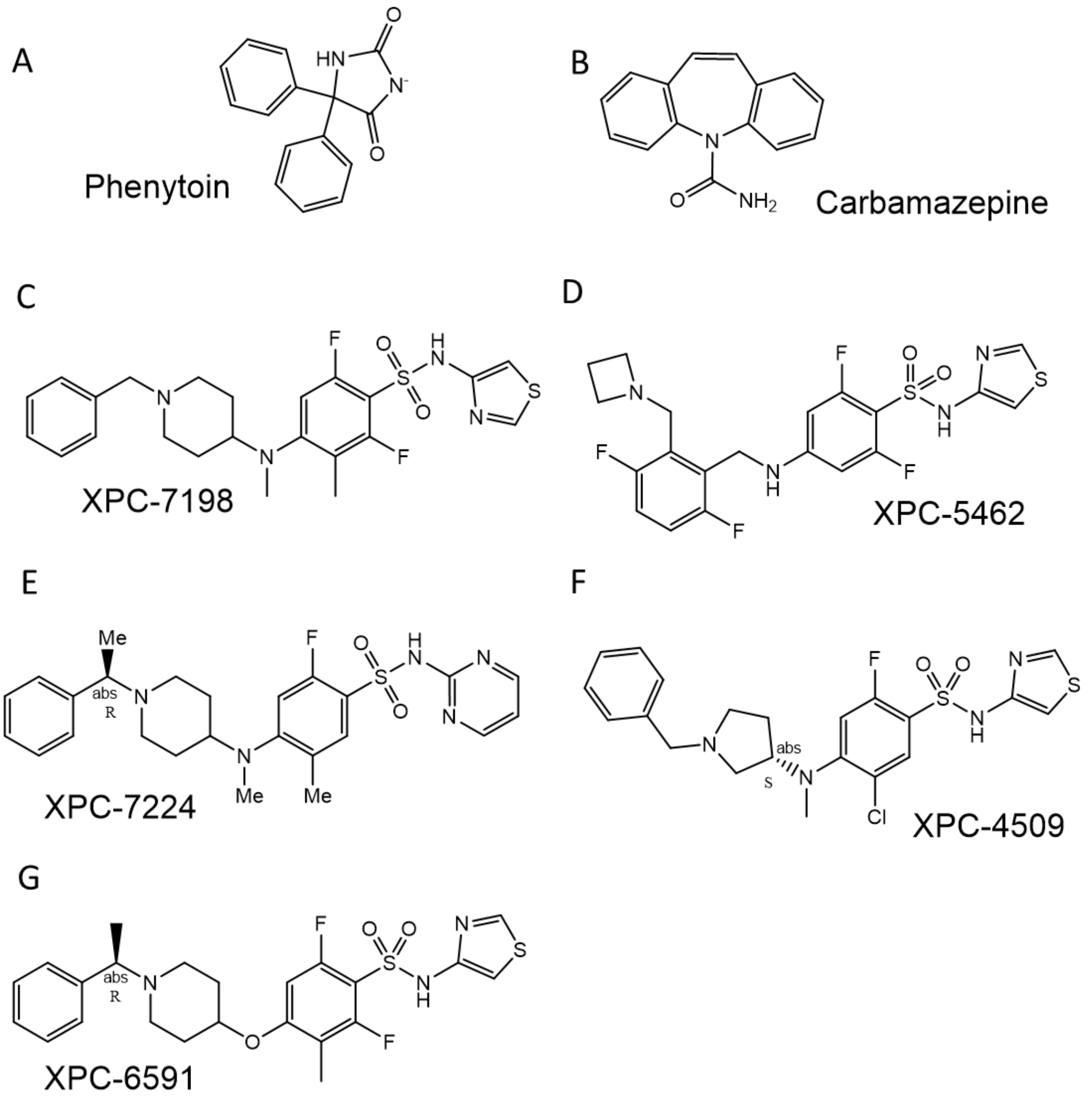


Phenytoin (A) and carbamazepine (B) are established anti-seizure medicines commonly used to treat patients. All XPC compounds (C-G) were created by, and patented by, Xenon Pharmaceuticals.

**Figure S2. Flux assay potency and selectivity.**

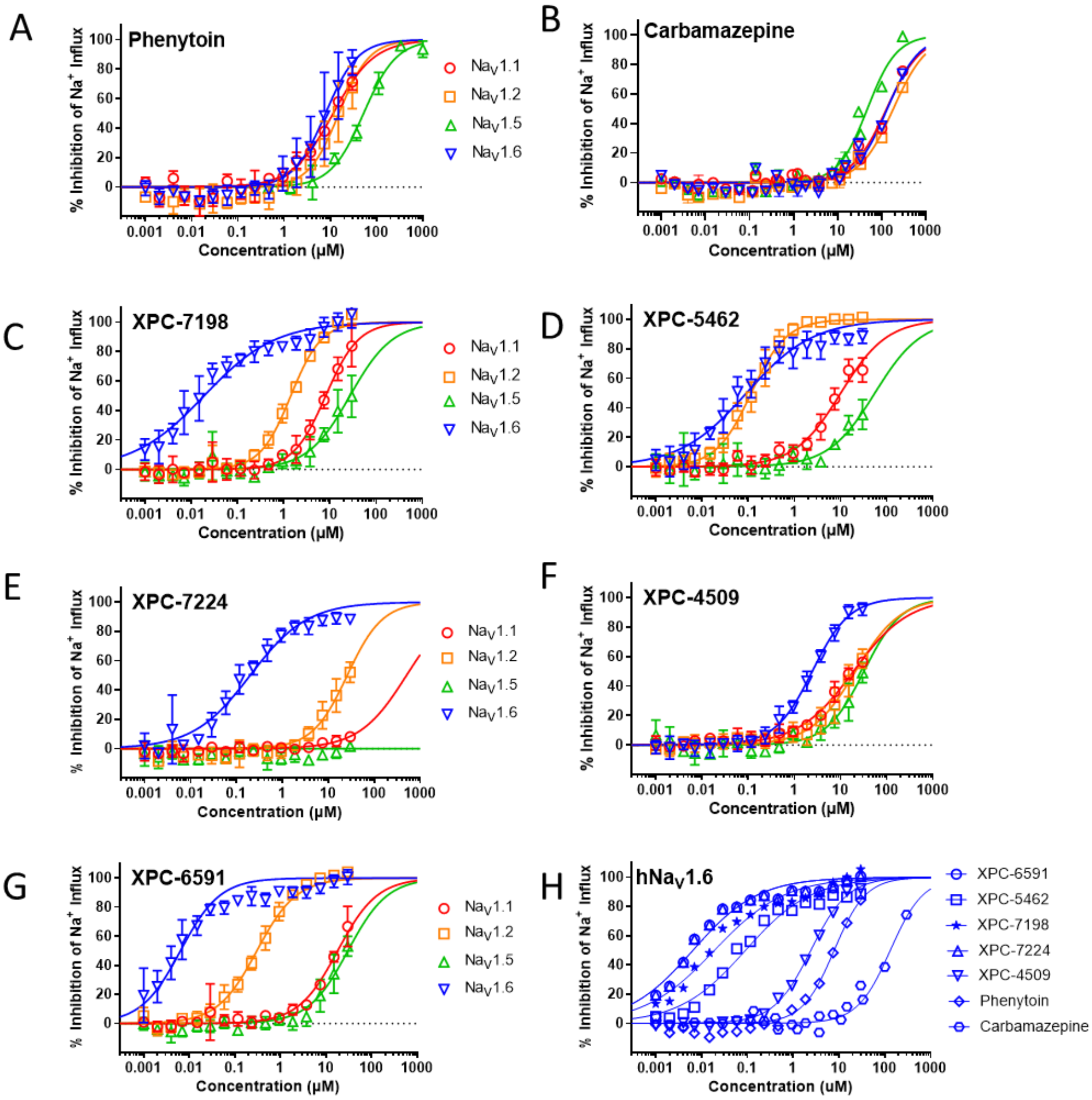


Potency of $Na_V$ inhibitor compounds for the adult human CNS and Cardiac $Na_V$ channels as determined by a fluorescent sodium influx assay. Phenytoin (A) and carbamazepine (B) have modest potency and little selectivity amongst isoforms. Panels C-G show potencies of 5 XPC compounds on human $Na_V$1.1, 1.2, 1.5, and 1.6. XPC-6591 is the most potent, followed by XPC-5462 > XPC-7198 > XPC-7224 > XPC 4509 > phenytoin > carbamazepine. Error bars indicate the standard deviations. XPC-5462 (D), XPC-7224 (E), XPC-4509 (F), and XPC-6591 (G) are novel $Na_V$ inhibitors with diverse potency and selectivity profiles that were created at Xenon Pharmaceuticals and licensed by Neurocrine Biosciences. (H) shows the relative potency of all compounds on $hNa_V$1.6.

**Figure S3: Potency on mouse Na$_V$1.6.**

A

| | | Na$_V$1.6 |
|---|---|---|
| **6591** | IC50 (µM) | 0.034 |
| | 95% CI | 0.023 to 0.049 |
| | N (cells) | 44 |
| **7198** | IC50 | 0.0021 |
| | 95% CI | 0.0049 to 0.016 |
| | N (cells) | 34 |
| **5462** | IC50 (µM) | 0.023 |
| | 95% CI | 0.016 to 0.033 |
| | N (cells) | 42 |
| **7224** | IC50 (µM) | 0.058 |
| | 95% CI | 0.13 to 0.43 |
| | N (cells) | 35 |
| **4509** | IC50 (µM) | 1.6 |
| | 95% CI | 1.1 to 2.3 |
| | N (cells) | 29 |
| **PHT** | IC50 (µM) | 11 |
| | 95% CI | 6.6 to 17 |
| | N (cells) | 52 |
| **CBZ** | IC50 (µM) | 17 |
| | 95% CI | 12 to 26 |
| | N (cells) | 36 |

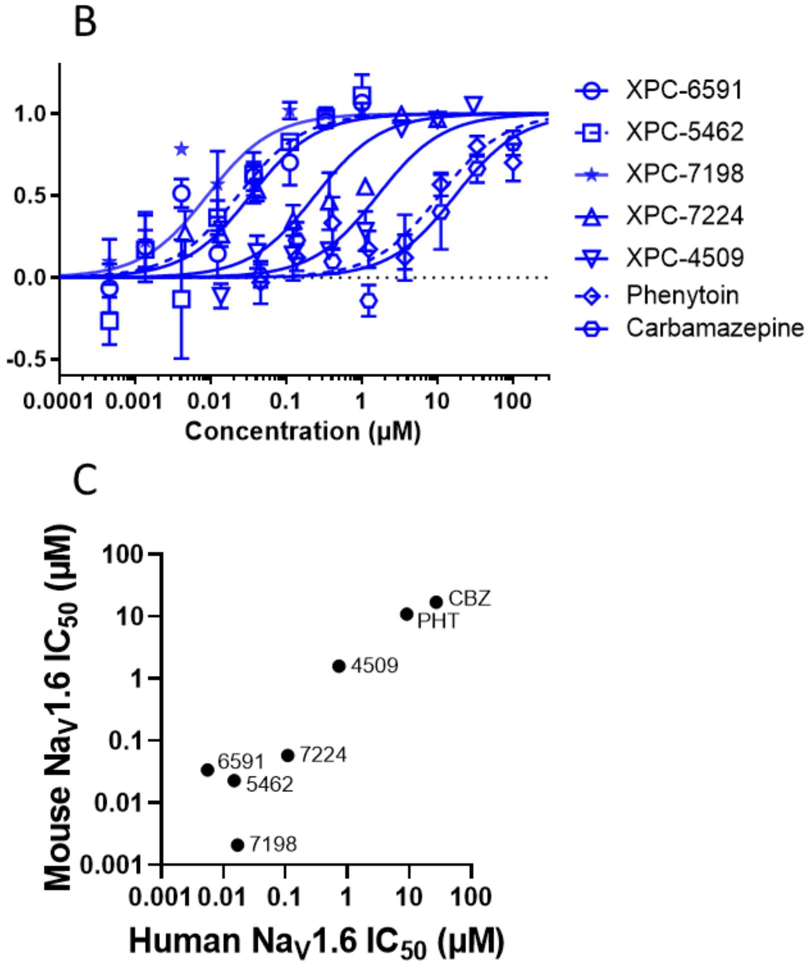


Potency of Na$_V$ inhibitor compounds for mouse Na$_V$1.6.IC$_{50}$ values in mice are closely aligned with those for human Na$_V$1.6.

**Figure S4: Scn8a statistics.**

A

| Dose (mg/kg) | P value | N |
|---|---|---|
| 1 | >0.9999 | 11 |
| 3 | 0.7094 | 11 |
| 10 | <0.0001 | 11 |
| 30 | <0.0001 | 11 |
| 56 | 0.0003 | 5 |

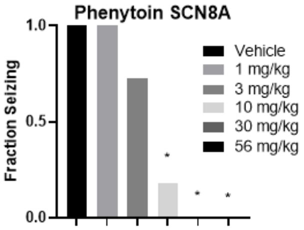


B

| Dose (mg/kg) | P value | N |
|---|---|---|
| 3 | 0.4644 | 12 |
| 10 | 0.5592 | 13 |
| 30 | 0.0009 | 5 |
| 100 | 0.0009 | 5 |

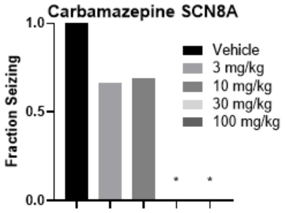


C

| Dose (mg/kg) | p value | N |
|---|---|---|
| 1 | 0.018 | 15 |
| 3 | <0.0001 | 19 |
| 10 | <0.0001 | 19 |
| 30 | <0.0001 | 14 |

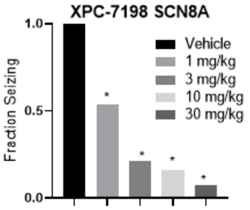


D

| Dose (mg/kg) | p value | N |
|---|---|---|
| 1 | >0.9999 | 6 |
| 3 | >0.9999 | 11 |
| 10 | 0.2462 | 11 |
| 30 | 0.0062 | 25 |
| 100 | <0.0001 | 16 |
| 200 | 0.0001 | 8 |
| 300 | <0.0001 | 9 |
| 400 | 0.0042 | 4 |

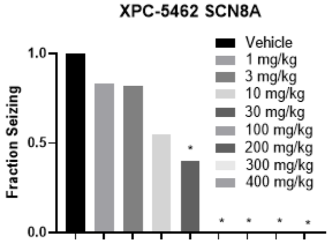


E

| Dose (mg/kg) | p value | N |
|---|---|---|
| 1 | >0.9999 | 7 |
| 3 | >0.9999 | 7 |
| 10 | 0.0129 | 13 |
| 30 | 0.0002 | 14 |
| 100 | <0.0001 | 8 |

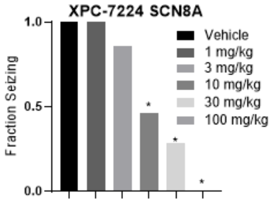


F

| Dose (mg/kg) | p value | N |
|---|---|---|
| 10 | >0.9999 | 6 |
| 30 | 0.1599 | 10 |
| 50 | 0.0676 | 5 |
| 100 | 0.0009 | 10 |
| 200 | <0.0001 | 12 |

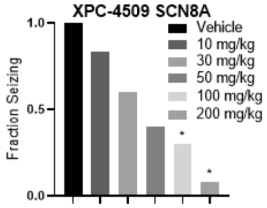


G

| Dose (mg/kg) | p value | N |
|---|---|---|
| 0.3 | >0.9999 | 6 |
| 1 | 0.1233 | 15 |
| 3 | 0.0002 | 14 |
| 10 | <0.0001 | 7 |

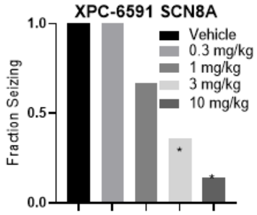


Number of animals, statistical significance and *P* values for the data in the left column of the tables that accompany each bar graph. Groups of animals treated with the same dose in different experimental runs were combined for purposes of analysis. Between-group differences were compared to vehicle response and were analyzed using a Kruskal-Wallis test followed by Dunn’s multiple comparisons test. *P*<.05 were considered significant.

**Figure S5: Mouse plasma and brain free fractions.**

A

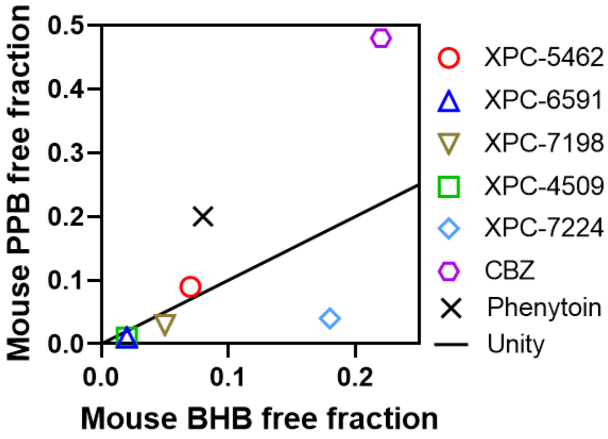


B

| | BHB free fraction | PPB free fraction |
|---|---|---|
| **XPC-5462** | 0.07 | 0.09 |
| **XPC-6591** | 0.02 | 0.01 |
| **XPC-4509** | 0.02 | 0.01 |
| **XPC-7198** | 0.05 | 0.03 |
| **XPC-7224** | 0.18 | 0.04 |
| **CBZ** | 0.22 | 0.48 |
| **Phenytoin** | 0.08 | 0.2 |

Compound free fractions as determined by equilibrium dialysis in mouse plasma or mouse brain homogenate.

**Figure S6: DC MES statistics.**

A

| Dose (mg/kg) | P value | N |
|---|---|---|
| 2 | 0.4093 | 13 |
| 5 | 0.1037 | 15 |
| 10 | <0.0001 | 7 |

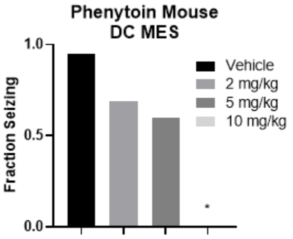


B

| Dose (mg/kg) | P value | N |
|---|---|---|
| 10 | 0.7862 | 14 |
| 30 | 0.0084 | 14 |
| 100 | 0.0006 | 7 |

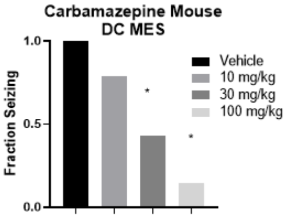


C

| Dose (mg/kg) | p value | N |
|---|---|---|
| 0.3 | >0.9999 | 7 |
| 1 | 0.2895 | 7 |
| 3 | 0.0595 | 7 |
| 10 | 0.0009 | 7 |
| 30 | 0.0086 | 7 |

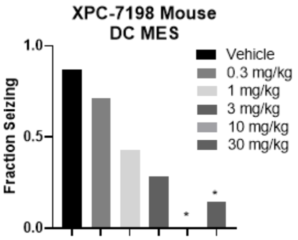


D

| Dose (mg/kg) | p value | N |
|---|---|---|
| 0.3 | >0.9999 | 7 |
| 2 | >0.9999 | 7 |
| 3 | >0.9999 | 8 |
| 5 | 0.5393 | 23 |
| 10 | 0.21 | 14 |
| 30 | 0.0009 | 23 |
| 100 | 0.0005 | 7 |

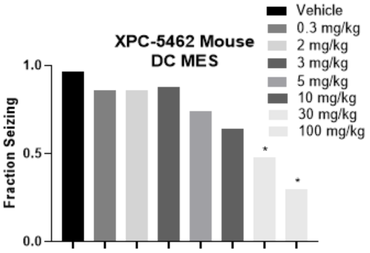


E

| Dose (mg/kg) | p value | N |
|---|---|---|
| 3 | 0.422 | 8 |
| 10 | 0.0096 | 14 |
| 30 | 0.0015 | 15 |
| 100 | 0.0096 | 4 |

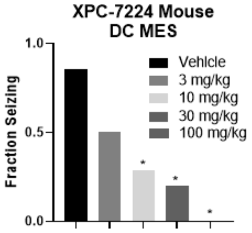


F

| Dose (mg/kg) | p value | N |
|---|---|---|
| 30 | >0.9999 | 8 |
| 56 | >0.9999 | 8 |
| 100 | 0.0538 | 8 |
| 200 | 0.0121 | 8 |

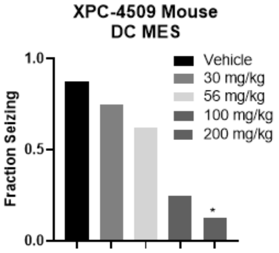


G

| Dose (mg/kg) | p value | N |
|---|---|---|
| 0.3 | >0.9999 | 16 |
| 1 | 0.4392 | 15 |
| 3 | 0.001 | 22 |
| 10 | 0.0011 | 8 |

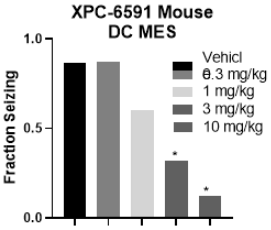


Number of animals, statistical significance and *P* values for the data in the left column of the tables that accompany each bar graph. Groups of animals treated with the same dose in different experimental runs were combined for purposes of analysis. Between-group differences were compared to vehicle response and were analyzed using a Kruskal-Wallis test followed by Dunn's multiple comparisons test. *P*<.05 were considered significant.
